# PBP1A and LdtJ support cell envelope homeostasis and impact selection of Colistin-resistance in *Acinetobacter baumannii*

**DOI:** 10.64898/2026.04.09.717377

**Authors:** Berenice Furlan, Sinjini Nandy, Hanling Gong, Michael B. Whalen, Nicolò Mattei, Roberto Jhonatan Olea-Ozuna, Daniela Vollmer, Waldemar Vollmer, Joseph M. Boll, Orietta Massidda

## Abstract

The multilayered cell envelope of *Acinetobacter baumannii* is an essential structure that maintains cellular integrity and protects the bacterial cell against external stresses and antibiotics. It consists of an inner membrane, a thin peptidoglycan (PG) layer and an asymmetric outer membrane (OM) enriched in lipooligosaccharide (LOS), whose lipid A moiety is the target of colistin, a last-resource antibiotic. Although lipid A is essential in most Gram-negatives, *A. baumannii* can survive without LOS through envelope remodeling, particularly in strains producing low levels of the bifunctional penicillin-binding protein PBP1A (encoded by *mrcA*) or in Δ*mrcA* mutants. Here, we identify a functional interplay between the LD-transpeptidase LdtJ, which generates 3-3 cross-links, and PBP1A, which catalyzes 4-3 transpeptidation during PG synthesis. We show that simultaneous inactivation of both enzymes severely affected growth, viability, morphology, and OM homeostasis. PG analyses revealed that the Δ*ldtJ* Δ*mrcA* mutants displays reduced overall cross-linkage and shorter glycan chains, producing a weakened sacculus. Co-immunoprecipitation demonstrated that PBP1A associates with LdtJ, supporting their coordinated activity at sites of PG synthesis. Notably, Δ*ldtJ* Δ*mrcA* mutants exhibited the highest recovery frequency of colistin-resistant, LOS-deficient variants compared with wild type or single mutants. Together, our findings demonstrate that coupling between 4-3 and 3-3 transpeptidation is critical for envelope stability in *A. baumannii* and highlight how disrupting this coordination favors the emergence of colistin resistance. This work identifies a conserved PG remodeling vulnerability that directly links PG integrity to the evolution of antibiotic resistance, offering a new conceptual framework for destabilizing the *A. baumannii* envelope.

**Importance:** The global rise of multidrug-resistant *Acinetobacter baumannii* represents an urgent clinical threat, largely driven by its extraordinary capacity to withstand cell envelope damages and escape last-resort antibiotics such as colistin. Although peptidoglycan synthesis and remodeling are known to influence outer membrane stability, how these processes are coordinated in *A. baumannii* has remained unclear. Here, we identify a crucial interplay between two critical envelope biogenesis and remodeling activities, 4-3 and 3-3 transpeptidations, mediated by the bifunctional PBP1A and the LD-transpeptidase LdtJ, respectively. Disrupting this coordination weakens the peptidoglycan, destabilizes the outer membrane, and increases the emergence frequency of colistin-resistant, LOS-deficient variants. These findings highlight a previously unrecognized vulnerability in the envelope homeostasis of *A. baumannii*, suggesting that simultaneously targeting DD and LD-transpeptidation could potentiate therapeutic strategies aimed at limiting antibiotic resistance development.

## Introduction

*Acinetobacter baumannii* is a Gram-negative opportunistic pathogen that has emerged as a major concern in healthcare settings due to its remarkable capacity to survive environmental stress and resist multiple classes of antibiotics (1), including last-resort agents such as colistin (2). This resilience largely relies on the structural and functional integrity of the cell envelope, a multilayered barrier essential for bacterial survival and defense against antimicrobial agents and environmental challenges (3).

The Gram-negative cell envelope consists of three principal components: the inner membrane (IM), the peptidoglycan (PG) layer, and the outer membrane (OM). The OM is particularly notable for its asymmetric lipid organization: the inner leaflet consists of phospholipids, whereas the outer leaflet is enriched with lipopolysaccharides (LPS), which together form a highly selective permeability barrier (3,4). In *A. baumannii*, this role is fulfilled by lipooligosaccharide (LOS), a truncated LPS variant lacking O-antigen polysaccharide (5). Despite this structural difference, LOS is essential for OM stability and protects the bacterium from toxic compounds and host immune factors.

Beneath the OM, the PG sacculus provides the cell with mechanical strength and osmotic stability (6). PG consists of alternating *N*-acetylglucosamine and *N*-acetylmuramic acid that are cross-linked by short peptides (7). In *Escherichia coli*, and many Gram-negative species, the predominant 4-3 cross-links are formed by the DD-transpeptidase activity of penicillin-binding proteins (PBPs), which link the fourth position D-alanine of a donor peptide to the third position *meso*-diaminopimelic acid (*m*DAP) of an acceptor peptide (8). An alternative reaction, the formation of 3-3 cross-links by LD-transpeptidases (LDTs), joins two *m*DAP residues and typically increases under conditions that inhibit classical PG synthesis or impose envelope stress (9–16).

In *A. baumannii*, 4-3 cross-links are similarly generated by PBPs, whereas the single LDT in this species, LdtJ, catalyzes 3-3 cross-links that contribute to cell envelope robustness (9,17–20). Although generally nonessential, LDTs are known to enhance resistance to β-lactam antibiotics (10–13,17,21,22), protect against lysozyme (9), and support envelope integrity (19) when OM assembly is impaired (15). LDT-mediated cross-linking often compensates for perturbations in PG architecture to maintain envelope function (17). However, the extent to which DD- and LD-transpeptidases coordinate PG synthesis and remodeling in *A. baumannii* remains poorly understood.

To address this gap, we analyzed single and double mutants lacking *mrcA* or *mrcB*, which encode class A PBPs PBP1A and PBP1B respectively, as well as *ldtJ*, in the multidrug-resistant (MDR) clinical isolate AB5075. Loss of *mrcA* causes pronounced cell elongation, consistent with a role for PBP1A in cell division (17,20), whereas deletion of *mrcB* produces minimal phenotypic defects (17). Simultaneous deletion of *mrcA* and *mrcB* is synthetically lethal (17), underscoring the essential and partially redundant functions. Previous work also proposed synthetic lethality between *mrcA* and *ldtJ*, suggesting that LdtJ-mediated 3-3 cross-linking becomes indispensable when PBP1A-dependent 4-3 cross-links are lost (17).

Here, we show that *ldtJ mrcA* double mutants are instead viable despite exhibiting severe growth defects, pronounced morphological abnormalities, and compromised envelope homeostasis. These phenotypes occur in both the clinical strain AB5075 and the laboratory strain ATCC 17978, indicating that the functional interplay between PBP1A and LdtJ is crucial, albeit not strictly essential, for *A. baumannii* cells.

We further examined how combined loss of *ldtJ* and *mrcA* influences the emergence of colistin-resistant, LOS-deficient (ColR LOS^-^) variants. Colistin exerts its bactericidal activity by binding the phosphate groups of lipid A and displacing divalent cations, thereby disrupting OM integrity (21,23). Although lipid A is essential in most Gram-negative bacteria, *A. baumannii* can survive without it through inactivating mutations in *lpxA*, *lpxC*, and *lpxD*, which block LOS biosynthesis (5,22). This LOS-deficient (LOS-) state imposes a substantial structural stress but can be tolerated when *mrcA* function is reduced or absent (22), highlighting the critical interdependence between PG and OM.

Collectively, our findings reveal that although *ldtJ* and *mrcA* are not synthetically lethal, their combined inactivation disrupts growth, cell morphology, and envelope integrity. These results uncover an underappreciated coordination between PG synthesis, maturation and remodeling pathways and OM homeostasis in *A. baumannii*.

## Results

### Deletion of *ldtJ* together with *mrcA*, but not with *mrcB*, impacts growth, viability and cell morphology

To investigate the functional interaction between LdtJ and PBP1A, we monitored growth of the *A. baumannii* MDR strain AB5075 and mutants lacking either or both corresponding genes. The Δ*ldtJ* Δ*mrcA* double mutant was generated in the Δ*ldtJ* background by removing first, *via* Cre recombination, the Tn26 tetracycline resistance cassette, followed by transformation with genomic DNA from the *ΔmrcA* mutant, as previously reported (24), and as detailed in the Materials and Methods. Whole-genome sequencing verified the absence of suppressor mutations, as shown in Fig. S1A and S1B. Likewise, a Δ*ldtJ* Δ*mrcB* double mutant was generated in strain AB5075 using the same strategy.

Deletion of *ldtJ* or *mrcA* alone in AB5075 did not show a significant impact on growth (Fig. 1A) or viability (Fig. 1B), in agreement with prior findings (17). In contrast, the Δ*ldtJ* Δ*mrcA* double mutant, albeit viable, exhibited impaired growth relative to wild type and both single mutants (Fig. 1A), coupled with a substantial reduction in viability (Fig. 1B). These findings indicate a critical role for LdtJ and PBP1A in maintaining PG homeostasis.

**Figure 1.**
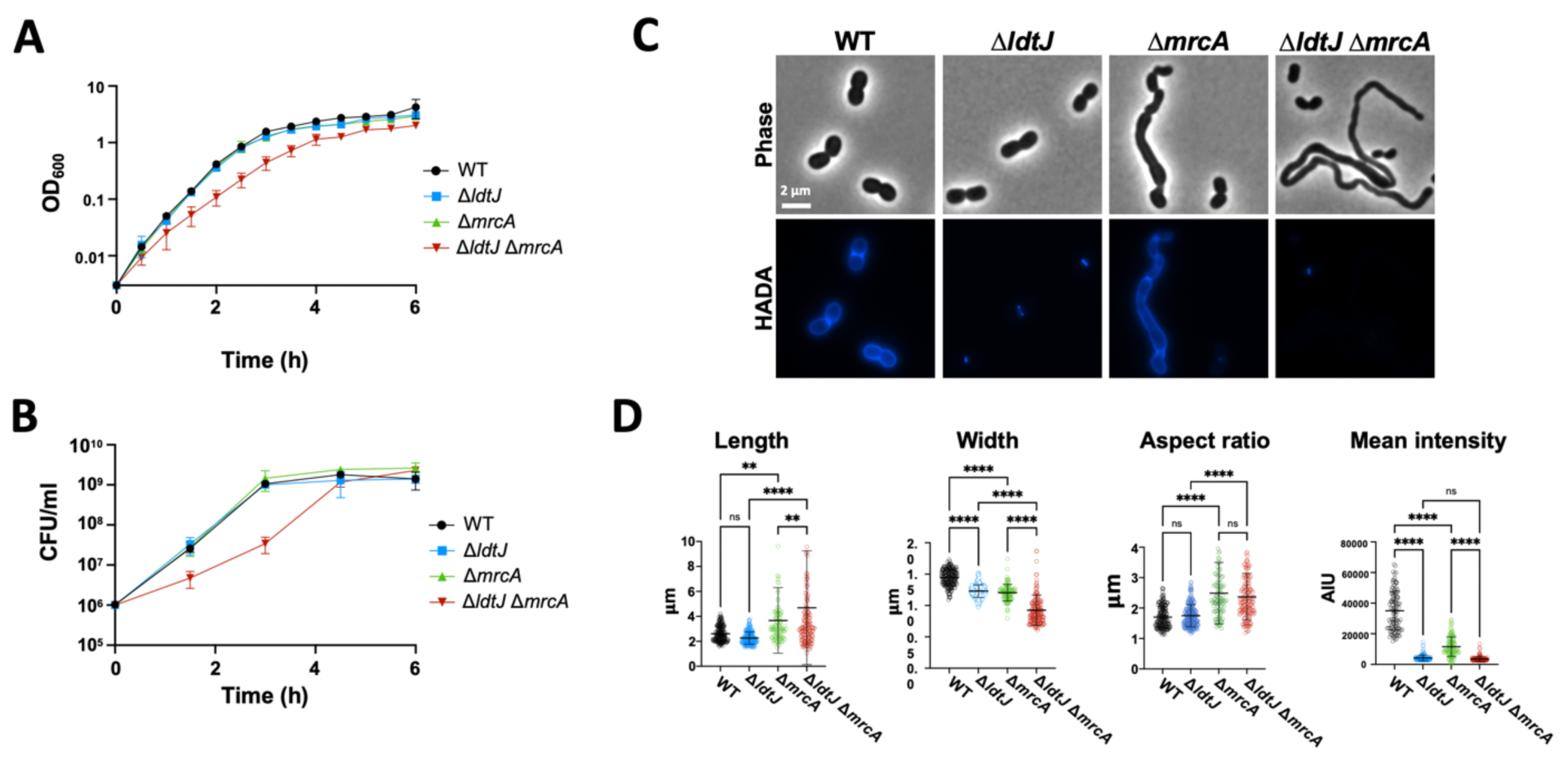
Characterization of the *A. baumannii* AB5075 Δ*ldtJ* Δ*mrcA* double mutant. **(A)** Growth (OD₆₀₀) and (B) viability (CFU/ml) of wild type (WT), Δ*ldtJ*, Δ*mrcA*, and Δ*ldtJ* Δ*mrcA* mutants in LB. Each experiment was independently performed in triplicate. (C) Phase-contrast and D-amino acid (HADA) fluorescence microscopy of the WT and mutants in logarithmic phase of growth. (D) Quantification of cell length, width, aspect ratio, and HADA fluorescence intensity (*n* >100 cells per strain), analyzed using ImageJ software. Statistical significance was determined using one-way ANOVA (\*\**P* < 0.01; \*\*\*\**P* < 0.0001; ns = not significant)

Given these growth and viability defects, we next examined cell morphology. The Δ*ldtJ* mutant displayed no appreciable differences compared to the wild type, whereas the Δ*mrcA* mutant exhibited an elongated, chain-like phenotype with multiple or incomplete septa (Fig. 1C), in line with the established role of *A. baumannii* PBP1A in cell division (17,20). The Δ*ldtJ* Δ*mrcA* double mutant showed severe morphological abnormalities, producing markedly elongated cells than the Δ*mrcA* single mutant (Fig. 1C). Quantitative measurements (*n* >100) confirmed that Δ*ldtJ* Δ*mrcA* cells were significantly longer and narrower compared to the wild type and single mutants (Fig. 1D).

Consistent with the compromised LD-transpeptidase activity, both the *ΔldtJ* single and the *ΔldtJ* Δ*mrcA* double mutant exhibited reduced incorporation of the fluorescent D-amino acid analogue HADA relative to wild type, leading to a significant decrease in fluorescence intensity (Fig. 1C and 1D). HADA is a non-canonical D-amino acid (NCDAA) predominantly incorporated into *A. baumannii* by LdtJ, thus specifically labeling sites of active PG synthesis (17).

In contrast, the Δ*ldtJ* Δ*mrcB* double mutant resembled the wild type and the respective single mutants with respect to growth and HADA incorporation, although these cells were significantly thinner than wild type and the single mutants (Fig. S2).

Collectively, these results indicate that simultaneous deletion of *ldtJ* and *mrcA*, but not *ldtJ* and *mrcB*, severely impacts the growth, viability, and morphology of *A. baumannii*. These data also underscore that, despite the semi-redundancy and synthetic lethality between PBP1A and PBP1B (17), the absence of PBP1A has a more profound impact for *A. baumannii*, particularly when coupled with loss of LdtJ.

We examined the Δ*ldtJ* Δ*mrcA* phenotype also in the laboratory strain ATCC 17978. In this background, deletion of *ldtJ* caused more pronounced defects than in AB5075, including a mild growth delay and shape abnormalities characterized by shorter, rounder cells (Fig. S3A, S3B and S3C; 17,19). As in AB5075, the ATCC 17978 Δ*mrcA* mutant displayed an elongated phenotype (17), and the Δ*ldtJ* Δ*mrcA* double mutant exhibited elongated cells and exacerbated morphological defects, (Fig. S3B and S3C).

Complementation analysis of the Δ*ldtJ* Δ*mrcA* double mutants with either *ldtJ* or *mrcA* provided further insight. Ectopic expression of *ldtJ* (pLdtJ) in Δ*ldtJ* Δ*mrcA* double mutants restored normal HADA incorporation but did not rescue cell shape in either AB5075 (Fig. S4A) or ATCC 17978 (Fig. S4B). Conversely, expression of *mrcA* (pPBP1A) restored cell morphology defects but did not restore HADA incorporation.

### Double inactivation of LdtJ and PBP1A also affects OM stability and integrity

PG provides mechanical support to the bacterium to prevent lysis from the internal turgor. In Gram-negative bacteria, the connection between PG and OM is an emerging theme (25). Although LdtJ and PBP1A are enzymes involved in PG synthesis, maturation and remodeling, their function may extend beyond PG assembly and editing, directly or indirectly. This relationship is grounded in the fact that the PG meshwork and the OM form a coordinated structural unit (25), where alterations in PG assembly can compromise OM anchoring and lipid organization, facilitating its destabilization under stress conditions.

To explore this intimate connection, we assessed the susceptibility of the AB5075 *ldtJ* and *mrcA* single and double mutants to agents challenging OM permeability, specifically rifampin (Fig 2A) at increasing concentrations and detergents, such as SDS or the combination of SDS and EDTA (Fig 2B). The assays revealed that, whereas the individual loss of either enzyme resulted in only small differences compared to the wild type, the simultaneous inactivation of *ldtJ* and *mrcA* caused up to a four-log (10000-fold) reduction in growth in the presence of rifampin, SDS and SDS/EDTA (Fig. 2A and 2B), suggesting increased OM permeability and loss of OM integrity.

**Figure 2.**
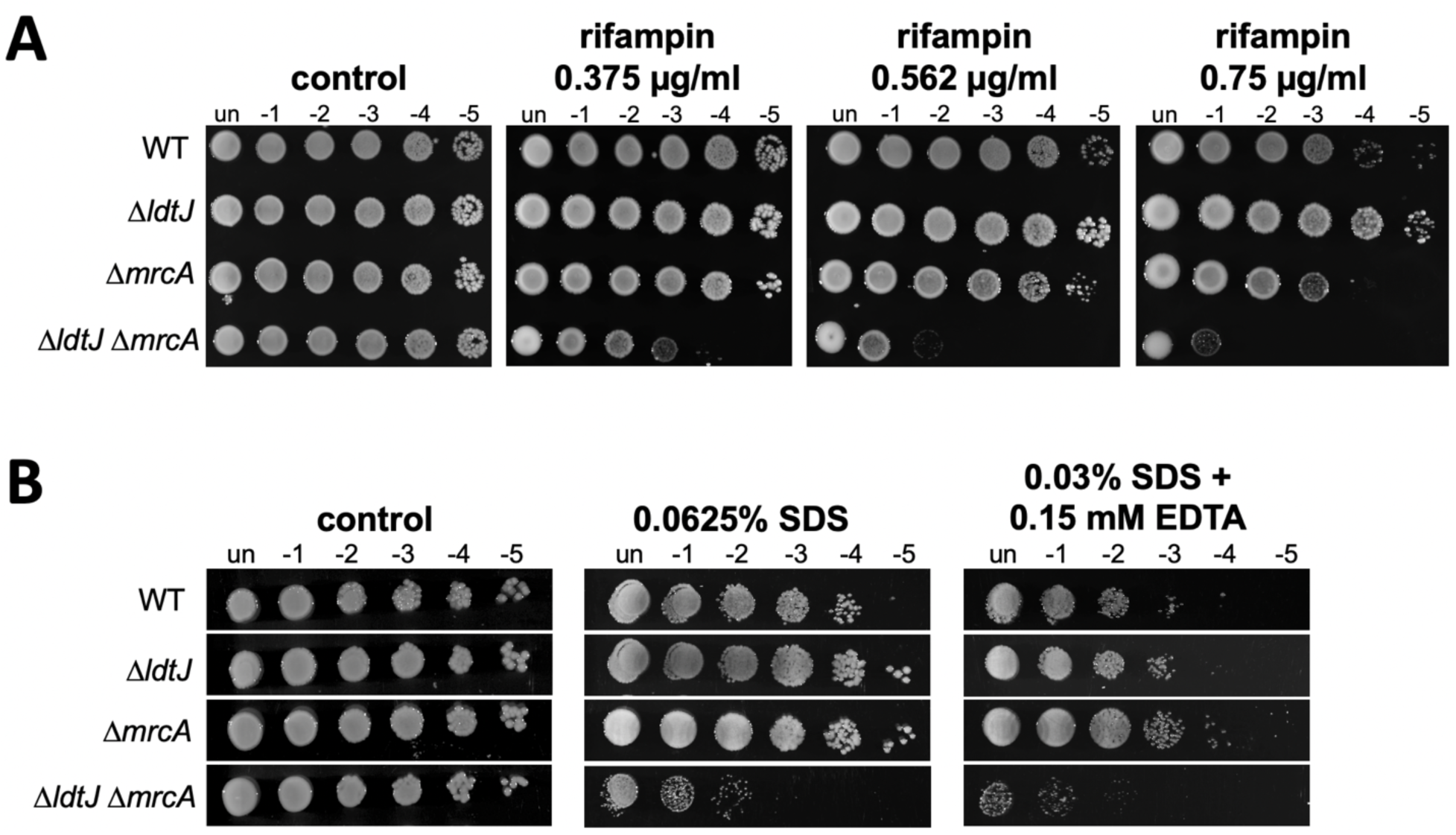
Defects of the Δ*ldtJ* Δ*mrcA* double mutant in OM permeability and integrity. Colony spot susceptibility assays of *A. baumannii* AB5075 WT, Δ*ldtJ*, Δ*mrcA* single mutants, and the Δ*ldtJ* Δ*mrcA* double mutant. Serial dilutions spotted on A) LB agar containing increasing concentrations of Rifampin and B) LB agar containing 0.0625% SDS or 0.03% SDS/0.15 mM EDTA.

To confirm that the observed phenotypes were directly caused by the loss of *ldtJ* and *mrcA*, we used the AB5075 Δ*ldtJ ΔmrcA* double mutant expressing ectopically each gene individually (Fig. S5). The Δ*ldtJ ΔmrcA* double mutant harboring the empty vector exhibited markedly reduced growth in the presence of rifampin (0.562 µg/ml), whereas ectopic expression of either LdtJ or

PBP1A partially or completely restored the ability to grow in the presence of rifampicin, respectively, rescuing the OM defects of the double mutant. These results reinforce the specific and non-redundant contributions of both enzymes in maintaining OM stability and integrity in *A. baumannii* and suggest that PBP1A plays a more prominent role than LdtJ in shaping PG structural organization, as its expression more effectively restores growth under rifampin stress conditions.

Together, these findings not only demonstrate a direct role of LdtJ and PBP1A in PG homeostasis but also highlight how PG maturation and remodeling critically influence OM structure and function. The exacerbated permeability defects in the Δ*ldtJ ΔmrcA* double mutant underscore the existence of parallel and partially redundant pathways that collaborate to uphold the bacterial barrier, reinforcing the concept of an integrated functional network between the PG and the OM in *A. baumannii*.

### Simultaneous inactivation of LdtJ and PBP1A redefines peptidoglycan architecture

To investigate the physiological consequences of the simultaneous inhibition of the transpeptidase activity of both LdtJ and PBP1A, we analyzed the PG composition of the AB5075 Δ*ldtJ* Δ*mrcA* double mutant in comparison with the single mutants during both logarithmic and stationary phases via HPLC and mass spectrometry (Fig. 3A and Fig. S6A).

**Figure 3.**
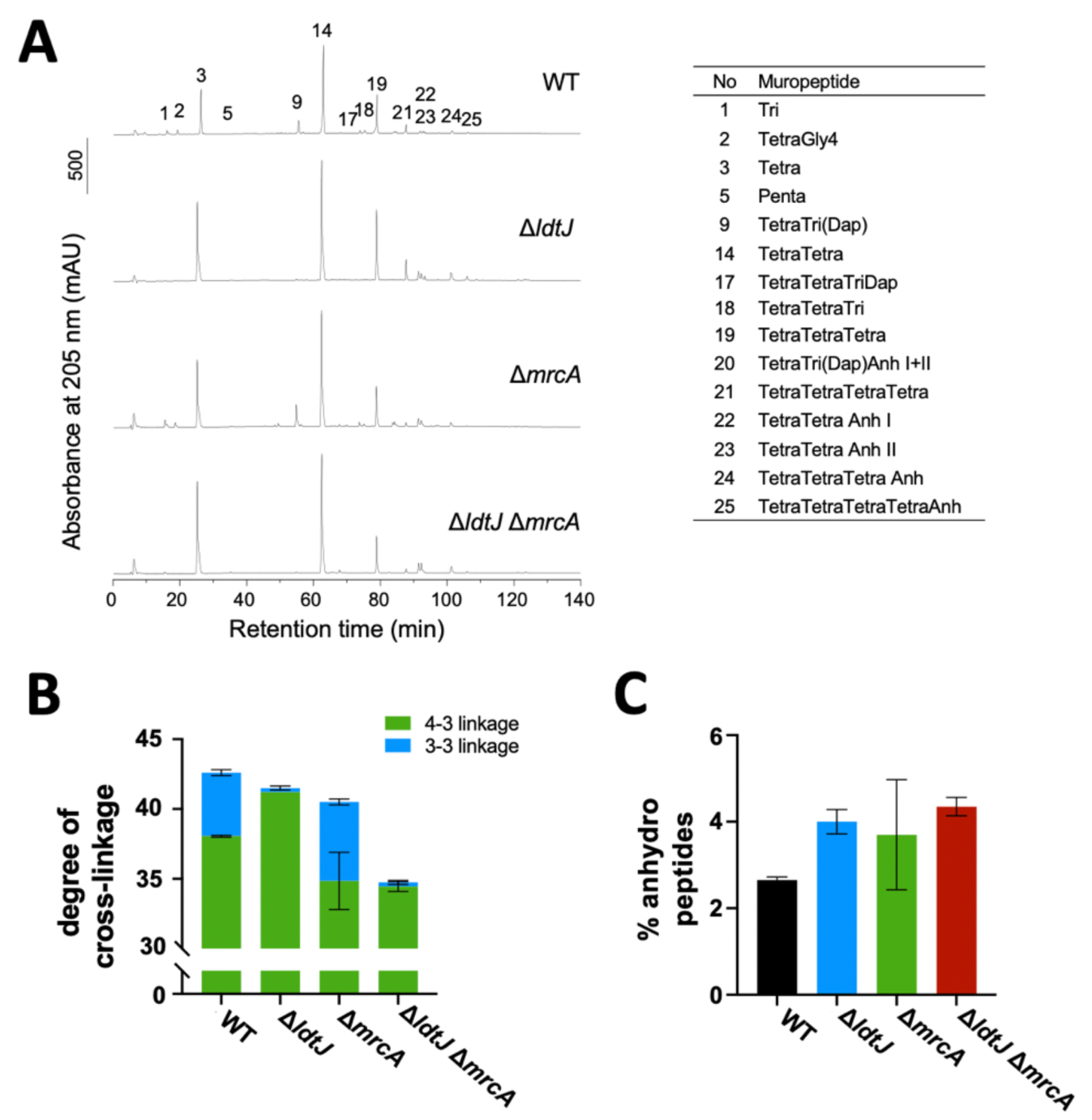
Peptidoglycan structure is altered in the *ΔldtJ ΔmrcA* double mutant. **(A)** Chromatograms showing PG composition of *A. baumannii* AB5075 WT, Δ*ldtJ*, Δ*mrcA* and the Δ*ldtJ* Δ*mrcA* mutants in logarithmic phase analyzed by HPLC. **(B)** Degree of 3-4 and 3-3 cross- links and **(C)** Percentage of total anhydro shortened chains. Error bars indicate variation of two biological replicates.

In logarithmic phase, Δ*ldtJ* mutant showed a near-complete loss of 3-3 cross-links as expected, confirming the pivotal role of LdtJ in catalyzing LD-transpeptidation also in the MDR strain AB5075 (Fig. 3A and 3B; 17,19). The Δ*mrcA* mutant slightly increased 3-3 cross-links (Fig. 3B), suggesting higher activity of LdtJ in the absence of PBP1A. Interestingly, the mutants differed in the levels of 4-3 cross-links in a similar way: as expected, the Δ*mrcA* mutant had decreased levels of 4-3 cross-links whilst 4-3 cross-links were increased in the Δ*ldtJ* mutant, suggesting that PBP1A activity was higher in the absence of LdtJ. Hence, cells appear to try compensating the loss of LdtJ or PBP1A by enhancing the activity of the remaining transpeptidase, resulting in altered contributions of 3-3 vs. 4-3 cross-links (Fig. S6B). Consistent with this, the double mutant lacking both *ldtJ* and *mrcA* showed a significant reduced overall cross-linkage (Fig. 3B). Another effect of the deletion of *ldtJ* was a strong reduction in tripeptides (Fig. 3A), presumably due to an inherent LD-carboxypeptidase activity of LdtJ (19). In stationary phase, the effects on LD 3-3 cross-links and tripeptides of Δ*ldtJ* were also observed, but the differences in 3-3 and 4-3 cross-links were less pronounced in all three mutants (Fig. S6 and Table S1). Anhydro-muropeptides levels, indicative of glycan strand length, increased in the Δ*ldtJ* mutant and to a higher extent also in the Δ*ldtJ* Δ*mrcA* double mutant in comparison to the wild type and the Δ*mrcA* mutant in exponential phase, indicating a more fragmented PG network (Fig. 3C).

Together, these results demonstrate that LdtJ and PBP1A make nonredundant contributions to overall PG composition and potentially compensate for the loss of the other in PG crosslinking during exponential growth and stationary phase. Their combined absence produces a structurally weakened sacculus, characterized by reduced 4-3 and 3-3 cross-links, and shorter glycan strands, indicating a compromised PG robustness. These combined defects provide a molecular explanation for the growth and viability impairment, altered morphology, increased OM permeability, and overall envelope instability observed in the Δ*ldtJ* Δ*mrcA* double mutant, where concurrent loss of LdtJ and PBP1A eliminates flexibility in PG synthesis, maturation and remodeling.

### Co-IP reveals a direct interaction between PBP1A and LdtJ in *A. baumannii*

Given the functional interplay observed between PBP1A and LdtJ, we next tested whether these proteins directly interact using co-immunoprecipitation (Co-IP). A C-terminal FLAG fusion protein, LdtJ-FLAG was expressed in both the ATCC 17978 wild type and Δ*mrcA* mutant (Fig. 4). Immunoprecipitation of LdtJ-FLAG with an anti-FLAG resin followed by immunoblotting revealed co-IP with PBP1A (Fig. 4), indicating that the two proteins associate *in vivo*, presumably within the same complex. Importantly, no signal was detected in the LdtJ-FLAG pull-down with the control protein RpoA, confirming interaction specificity. Immunoblots with the whole-cell lysates were included as controls as detected by specific antibodies and Anti-FLAG antibody (Fig. 5A) To further validate this association, PBP1A-FLAG was also expressed in the wild type and Δ*ldtJ* mutant (Fig 4) and subjected to Co-IP (Fig 4). LdtJ was consistently Co-IP’d with PBP1A-FLAG, whereas non-specific proteins were absent. Together, these reciprocal Co-IP assays provide evidence that PBP1A and LdtJ are part of the same protein complex *in vivo*.

**Figure 4.**
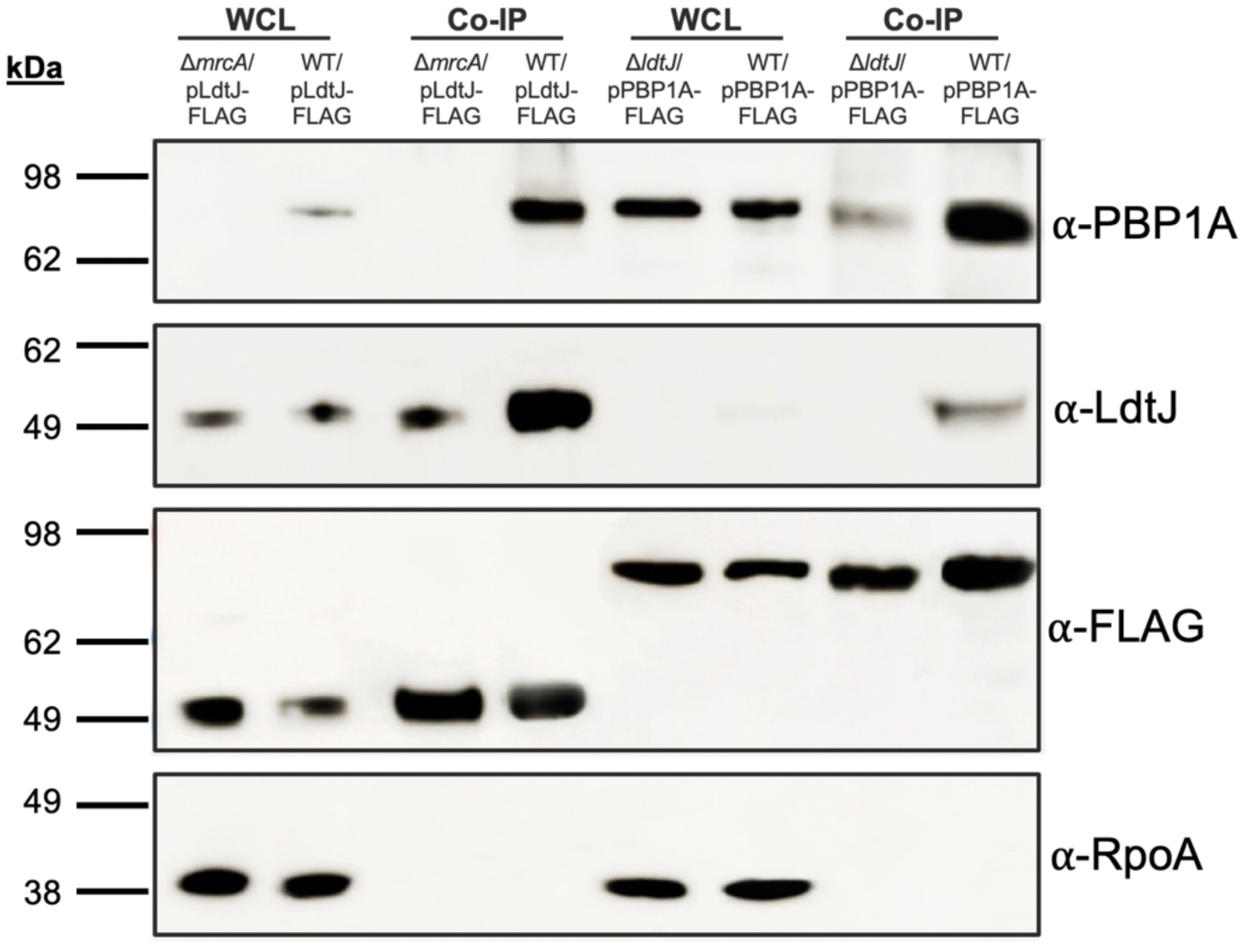
PBP1A directly interacts with LdtJ. Immunoblotting of the whole-cell lysates (WCL) controls and co-immunoprecipitation (Co-IP) fractions from *A. baumannii* ATCC 17978 wild type and Δ*mrcA* mutants expressing LdtJ-FLAG (1 and 2). LdtJ-FLAG was immunoprecipitated using anti-FLAG resin. Immunoblotting with anti-PBP1A antibody detected PBP1A specifically in LdtJ-FLAG pull-downs, indicating an interaction. Reciprocal Co-IP of PBP1A-FLAG expressed in WT or Δ*ldtJ* similarly co-precipitated LdtJ, confirming the association (3 and 4). PBP1A and LdtJ were detected with specific anti-PBP1A and anti-LdtJ antisera, LdtJ-FLAG and PBP1A-FLAG were detected using anti-FLAG monoclonal antibodies. RpoA (RNA polymerase α-subunit), also detected with a specific antiserum, was used as a cytoplasmic contamination control, confirming the specificity of the Co-IP fractions. PBP1A is 94.7 kDa; LdtJ is 44.5 kDa; FLAG tag is 1 kDa; RpoA is 37.6 kDa.

**Figure 5.**
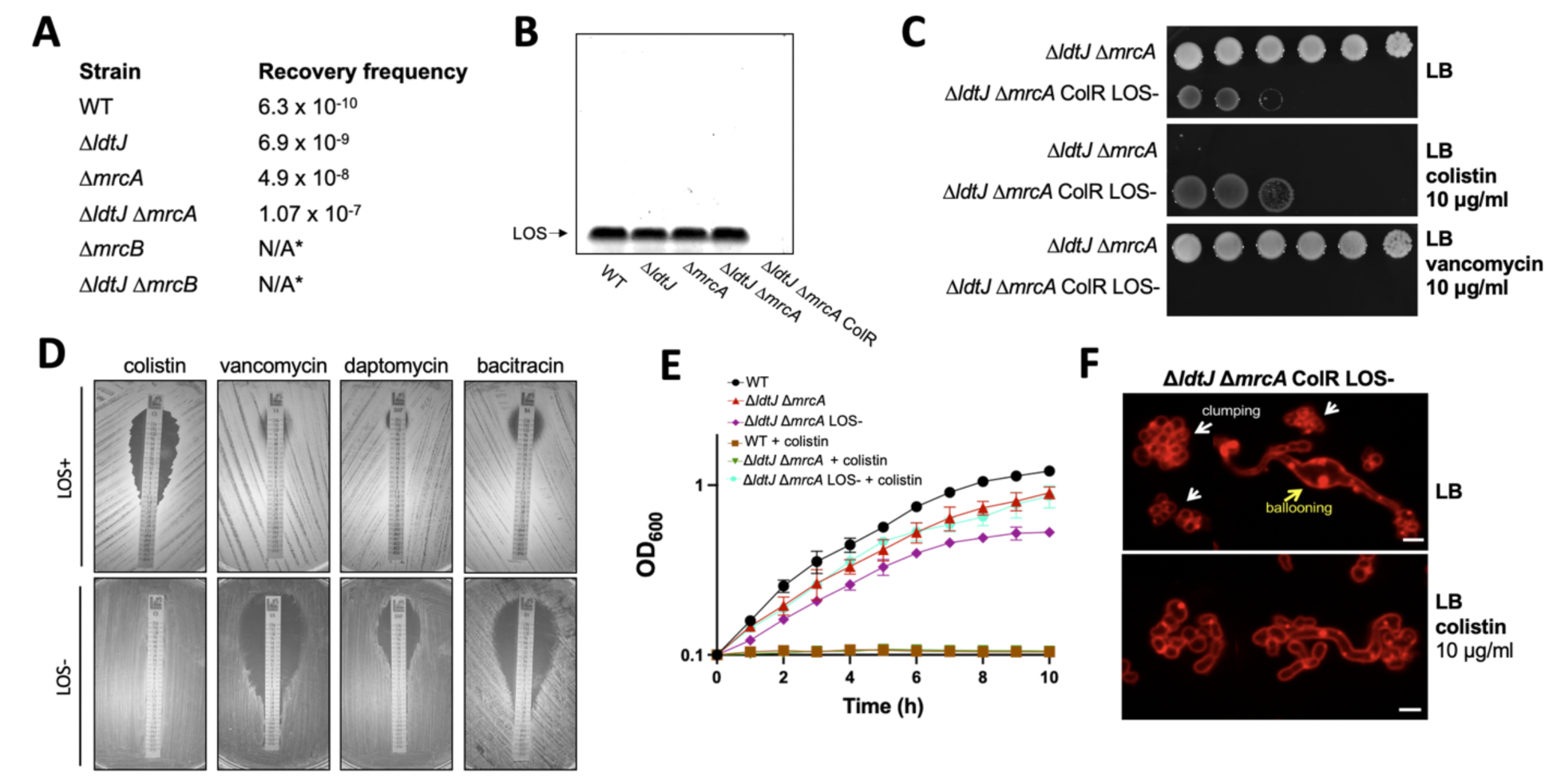
Isolation and characterization of the ColR LOS⁻ variants in the Δ*ldtJ* Δ*mrcA* double mutant. **(A)** Recovery frequencies of Col-R colonies from *A. baumannii* 5075 WT, Δ*ldtJ*, Δ*mrcA,* Δ*mrcB* single mutants and Δ*ldtJ* Δ*mrcA* and Δ*ldtJ* Δ*mrcB* double mutants. Recovery frequency of Δ*ldtJ* Δ*mrcA* and Δ*ldtJ* Δ*mrcB* Col-R colonies are the average values of two independent isolates for each mutant. *N/A: no Col-R LOS- isolates recovered. **(B)** LOS staining of the *A. baumannii* 5075 WT, Δ*ldtJ*, Δ*mrcA* single and the Δ*ldtJ* Δ*mrcA* COL-S and COL-R double mutants. **(C)** Colony spot assay of the Δ*ldtJ* Δ*mrcA* Col-S LOS+ parent and Col-R LOS⁻ derivative on LB agar and LB agar supplemented with Colistin (10 µg/ml) or Vancomycin (10 µg/ml). **(D)** Antibiotic susceptibility profile of the AB5075 Δ*ldtJ* Δ*mrcA* ColS LOS+ parent and ColR LOS⁻ derivative for Colistin and selected OM low-permeability antibiotics. **(E)** Growth curves of AB5075 WT, the Δ*ldtJ* Δ*mrcA* Col-S LOS+ parent and the Δ*ldtJ* Δ*mrcA*.

The identification of this interaction provides a possible mechanistic explanation for the synergistic phenotypes observed in the Δ*ldtJ* Δ*mrcA* double mutant. Since PBP1A synthesizes classical 4-3 cross-linking and LdtJ catalyzes the alternative 3-3 linkages, their physical interaction suggests that both enzymes may colocalize at sites of active PG assembly. Such spatial coordination would enable a direct coupling, or alternatively competition, between 4-3 and 3-3 cross-linking activities, thereby optimizing PG robustness during growth and division under stress conditions.

### Additive recovery of LOS-deficient, colistin-Resistant variants in the Δ*ldtJ* Δ*mrcA* mutant

The ability to generate colistin-resistant variants through loss of LOS varies among *A. baumannii* strains (17,22). While strains such as AB5075 and ATCC 19606 can naturally develop colistin resistance, strain ATCC 17978 is dependent on LOS for viability unless mutations, such as *mrcA* inactivation (17,22), are present to produce a permissive state. In this context, it is critical to understand how the combined inactivation of *ldtJ* and *mrcA* influences the frequency and phenotype of colistin-resistant variants in AB5075.

Determination of the LOS^-^ recovery frequency revealed low frequency of ColR LOS⁻ variants in the AB5075 wild type of 6.3 × 10⁻¹⁰. Single Δ*ldtJ* and Δ*mrcA* mutants showed increased recovery frequencies to 6.9 × 10⁻⁹ and 4.9 × 10⁻⁸, respectively. The Δ*ldtJ* Δ*mrcA* double mutant, moreover, exhibited the highest recovery frequency of 1.07 × 10⁻^7^, reflecting an effect that substantially promotes the selection of ColR LOS⁻ variants (Fig. 5A). In contrast, the single Δ*mrcB* mutant, and the Δ*ldtJ* Δ*mrcB* double mutant, failed to generate ColR LOS⁻ variants under the tested conditions (Fig. 5A). Similar experiments in the ATCC 17978 genetic background, where spontaneous selection of ColR LOS⁻ variants is inaccessible (22), showed no recovery of ColR variants in the wild type or in the Δ*ldtJ* mutant, as previously shown (17,22). The Δ*mrcA* mutant exhibited a frequency of 4.4 × 10⁻⁹ and the Δ*ldtJ* Δ*mrcA* double mutant showed a relative increase in recovery frequency of 7.4 × 10⁻⁸, supporting the conclusion that the combined inactivation of *mrcA* and *ldtJ* favors the selection of ColR LOS^-^ variants across different *A. baumannii* genetic backgrounds.

To test the lack of LOS in the isolated ColR variants generated by the AB5075 Δ*ldtJ* Δ*mrcA* mutant, proteinase K-treated cell lysates were analyzed for relative LOS levels by SDS-PAGE analysis. The wild type, the Δ*ldtJ* and Δ*mrcA* single mutants exhibited comparable LOS levels, as did the Δ*ldtJ* Δ*mrcA* parental strain (Fig. 5B). However, the Δ*ldtJ* Δ*mrcA* ColR variant lacked detectable LOS, confirming a LOS- phenotype (Fig. 5B). Moreover, the AB5075 Δ*ldtJ* Δ*mrcA* ColR LOS- variant displayed reduced viability (CFU/ml) on LB agar compared to the Δ*ldtJ* Δ*mrcA* parental strain (Fig. 3C, upper panel), suggesting a fitness cost associated with LOS deficiency, as has been reported with LOS- strains (26,27). Yet, in the presence of colistin (10 µg/ml), only Δ*ldtJ* Δ*mrcA* ColR LOS⁻ variant grew while the Δ*ldtJ* Δ*mrcA* parental strain did not grow (Fig 3C, middle panel). On the contrary, the presence of vancomycin (10 µg/ml) completely inhibited the growth of the Δ*ldtJ* Δ*mrcA* ColR LOS⁻ variant but not of the parental strain, highlighting altered OM permeability linked to LOS deficiency (Fig. 5C, bottom panel). The Minimum Inhibitory Concentrations (MICs), determined by E-test, demonstrated that the Δ*ldtJ* Δ*mrcA* parental strain was susceptible to colistin but resistant to vancomycin, bacitracin, and daptomycin, in line with previous reports on LOS- *A. baumannii* strains (22,47). Conversely, the Δ*ldtJ* Δ*mrcA* ColR LOS_⁻_ variant was resistant to colistin but became susceptible to the other antibiotics (Fig. 5D), presumably reflecting significant changes in OM permeability and consequent decreased MICs associated with LOS deficiency.

Growth curve analyses (OD_600_) corroborated these observations: the wild type and the Δ*ldtJ* Δ*mrcA* ColS LOS+ parental strain grew well in LB broth but was inhibited by colistin, whereas the ColR LOS_⁻_ variant not only grew in the presence of colistin but also exhibited enhanced growth rates compared to antibiotic-free conditions (Fig. 5E), suggesting an advantage under colistin pressure. We further characterized the Δ*ldtJ* Δ*mrcA* ColR LOS- variants by fluorescence microscopy using FM5-95 membrane staining. The Δ*ldtJ* Δ*mrcA* ColR LOS- variant exhibited cell “clumping” and “ballooning” when grown in LB without colistin; these phenotypes diminished notably upon exposure to colistin (Fig. 5F), which presumably stabilizes the remodeled OM during antibiotic stress (26, 27).

Together, these data demonstrate that simultaneous inactivation of *ldtJ* and *mrcA* increases the recovery frequency of ColR LOS- variants providing new insights into the compensatory mechanisms that *A. baumannii* employ to survive antimicrobial stress.

## Discussion

The coordination between PG synthesis and remodeling and OM integrity represents a fundamental aspect of envelope homeostasis in Gram-negative bacteria. In *A. baumannii*, however, this interplay is yet largely unexplored despite its critical role in antibiotic resistance (17,28,29) and virulence (30,31).

Our results demonstrate that the LD-transpeptidase LdtJ and the septal bifunctional class A PBP1A act in a coordinated manner to preserve envelope stability and integrity. Contrary to previous hypotheses (17), the simultaneous inactivation of LdtJ and PBP1A is not lethal but produces severe phenotypic and fitness defects. While the coordinated transpeptidation appears conserved, strain dependent effects were also observed. Inactivation of LdtJ alone in AB5075 does not induce obvious defects in cell growth, viability, or morphology (Fig. 1A and Fig. 1B), while in the 17978 Δ*ldtJ* mutant morphology is perturbed (Fig. S3). However, in both strains Δ*ldtJ* Δ*mrcA* double mutants showed severe defects characterized by reduced growth rate and viability, elongated and multiseptated cells with incomplete septa, and compromised OM homeostasis. Our PG analysis highlights that the 3-3 cross-linking formed by LdtJ and the 4-3 crosslinks synthetized by PBP1A represent parallel PG biosynthetic, maturation and remodeling activities that ensure a robust PG network and, together, confer the stability and the structural support to the cell envelope.

While the model organism *E. coli* has six LDTs, of which the LdtE generates most 3-3 cross-links in stationary phase (8,32) and LdtD becomes essential in complex with PBP1B and the DD-carboxypeptidase PBP6a to support the PG structure when the transport of the LPS to the OM is compromised (15), *A. baumannii* can only rely on LdtJ for the synthesis of the 3-3 cross-links. *E. coli* LdtD has been proposed to work in concert with the PBP1B, the major class A PG synthase involved in cell division, forming a “PG repairing” complex with PBP6a, which trims the pentapeptides to increase the pool of tetrapeptides to form the 3-3 cross-links. This machinery would reinforce the cell wall structure repairing the PG meshwork when the OM is compromised, providing an important model to demonstrate the intimate connection between PG remodeling and OM integrity (15). According to our data, it is reasonable to propose a model where LdtJ and PBP1A act within the same functional complex to optimize PG synthesis, maturation and remodeling. When one enzyme is inactivated, the other can partially compensate; however, the simultaneous loss of both enzymes results in severe defects in PG structure and integrity.

In *E. coli*, PBP1A and PBP1B are semi-redundant enzymes that can compensate for each other, and LDTs are generally dispensable under normal growth conditions (33,34). In contrast, our results indicate that when LD-transpeptidation is lost in *A. baumannii*, PBP1B cannot compensate for the loss of PBP1A. Furthermore, double inactivation of LdtJ and PBP1A, unlike inactivation of LdtJ and PBP1B, severely compromises the envelope integrity. This divergence suggests that, unlike *E. coli*, *A. baumannii* may have a reduced capacity to compensate for the loss of key PG biosynthetic enzymes, likely due to the limited functional redundancy in its genome.

The effects of the double LdtJ and PBP1A inactivation emerged also from our PG analysis. Consistent with LdtJ being the sole LDT in *A. baumannii* and with previous studies in the ATCC 17978 background (17,19), the Δ*ldtJ* mutant exhibited almost complete loss of 3-3 linkages and a decreased pool of total tripeptides fraction in comparison to the wild type. However, no differences in the incorporation of NCDAAs, other than D-Ala, were observed in the AB5075 Δ*ldtJ* mutant, differently from what reported in the ATCC 17978 Δ*ldtJ* (17,19) and in other Gram-negative bacteria like *E. coli* and *Vibrio cholerae* (35,36). This can be explained by the fact that also the AB5075 wild type doesn’t show incorporation of NCDAAs.

These strain-dependent differences between *A. baumannii* different genetic backgrounds could possibly reflect variations in the pool of LdtJ tetrapeptides substrates or acceptors and may provide an explanation for the different morphological phenotypes observed in the AB5075 ω*ldtJ* (wild type-like) and ATCC 17978 ω*ldtJ* (rounder) mutants.

In contrast, the Δ*mrcA* mutant consistently accumulated 3-3 cross-links across both genetic backgrounds compared to the wild type. This feature, interpreted as a possible requirement to substitute the 4-3 cross-links with more 3-3 cross-links in the absence of PBP1A (17), can now be more plausibly considered an indirect consequence of increased availability of tetrapeptide substrates for LdtJ, rather than a direct compensatory mechanism to reinforce PG structure. Strikingly, the Δ*ldtJ* Δ*mrcA* double mutant nearly abolished 3-3 links but also significantly reduced overall cross-linkage, with a drop in the average glycan chain length, yielding a sacculus that is less cross-linked and structurally weakened. These changes provide a direct molecular explanation for the reported growth, viability, morphological defects, increased OM permeability and instability observed in the Δ*ldtJ* Δ*mrcA* double mutant. In *E. coli*, LDTs, such as LdtD and LdtE, are known to become particularly important under β-lactam-induced stress or in strains with perturbed PG synthesis (13). Our results suggest that a similar stress-compensatory role for LD-transpeptidation also exists in *A. baumannii*, but that its interplay with PBP1A is more critical for maintaining basal envelope integrity.

The envelope defects in Δ*ldtJ* Δ*mrcA* extend to the OM permeability and integrity that was probed by testing its susceptibility to the hydrophobic antibiotic rifampin and the detergent SDS alone or combined with the chelator EDTA. The Δ*ldtJ* Δ*mrcA* double mutant showed a synergistic increase in susceptibility to rifampin, and to detergent/chelator combinations (SDS and SDS/EDTA), as it is more sensitive than the single mutants at all the probed concentrations and conditions. These defects were rescued by ectopic expression of either *ldtJ* or *mrcA*. Because the PG and OM form a mechanically coupled unit (37), a reduction in PG cross-linking likely impacts the OM structure altering its permeability and integrity properties. We propose that the combined loss of 3-3 cross-links and the reduced efficiency of 4-3 cross-linking disrupts the tension balance across the envelope, lowering the threshold for OM failure under chemical stress.

We also show that PBP1A physically interacts with LdtJ *in vivo*. The immunoprecipitation of PBP1A-FLAG recovered LdtJ, and the reciprocal co-IP of LdtJ-FLAG recovered PBP1A, supporting the idea that these enzymes operate in the same molecular complex. Such direct interaction may facilitate the spatial coordination of 4-3 and 3-3 cross-linking activities at the septum, ensuring a robust PG synthesis and remodeling. In *E. coli*, LdtD has been shown to physically interact with the class A septal PBP1B forming a complex in cells with OM assembly defects by fortifying the sacculus and preventing cell lysis (15). Beyond its role in supporting the cell envelope homeostasis, the association between LdtD and PBP1B and PBP5 has been proposed to strategically localize LdtD and facilitate its access to PBP1B and PBP5 substrates, mediating in this way β-lactam resistance (38).

Finally, a striking evolutionary consequence of disrupting this coordination was the increased selection of ColR LOS- variants in the Δ*ldtJ* Δ*mrcA* mutant. These findings suggest that the destabilization of the PG network can drive adaptive remodeling in the OM. In AB5075, the recovery frequency of ColR LOS- increased from wild type to the single mutants and was highest in the double mutant. A similar additive trend, though at lower absolute frequencies, was observed in the ATCC 17978. The Δ*ldtJ* Δ*mrcA* ColR variants lacked detectable LOS by gel staining, became susceptible to vancomycin and other antibiotics with poor OM penetration. The Δ*ldtJ* Δ*mrcA* ColR LOS- variants also exhibited morphology abnormalities, which were exacerbated when grown in drug-free conditions. Although slower growth rates and altered cell shape in *A. baumannii* LOS- have been reported (39), the mechanism by which collapse of the PG-OM system supports LOS deficiency while maintaining lipid bilayer integrity remains unclear. From a therapeutic perspective, the fitness costs observed in the Δ*ldtJ* Δ*mrcA* ColR LOS- variants suggest that the simultaneous inhibition of DD- and LD-transpeptidation combined with OM-permeabilizing adjuvants could potentiate envelope destabilization. The LdtJ-PBP1A system may therefore represent a druggable target in MDR *A. baumannii* clinical isolates.

Together, this work underscores that LdtJ and PBP1A form a functionally integrated system that coordinates PG synthesis, maturation and remodeling with OM stability and integrity. Balanced synthesis of 4-3 and 3-3 cross-links at the sites of active PG synthesis ensures mechanical strength and flexibility of the sacculus, thus contributing to the preservation of OM barrier function. Disruption of this equilibrium, however, paves the way for adaptive mechanisms that allow the development of colistin resistance via loss of LOS.

## Material and Methods

### Bacterial strains and growth conditions

All bacterial strains and plasmids used in this study are listed in Table S2, and the oligonucleotides are provided in Table S3. *A. baumannii* and *E. coli* were routinely grown aerobically on Luria-Bertani agar or Lysogeny Broth (LB) at 37°C from frozen stocks preserved at 30% glycerol at - 80°C. For selection, antibiotics were added to the growth media, when necessary, at the following final concentrations: tetracycline (5 µg/ml for AB5075 or 10 µg/ml for ATCC 17978) (40), hygromycin (250 µg/ml for AB5075) (40), kanamycin (25 µg/ml), colistin (10 µg/m), and vancomycin (10 µg/ml). Diaminopimelic acid (DAP) was added at 600 μM to support the growth of *E. coli* DAP^-^ donor strain. IPTG (isopropyl β-D-1-thiogalactopyranoside) was added at varying concentrations to induce gene expression as indicated.

### Construction of AB5075 *A. baumannii* double mutants and ectopic expression of *ldtJ* and *mrcA*

Isogenic mutants with transposon insertion were obtained from the *A. baumannii* AB5075 ordered mutant library of the Laboratory of Colin Manoil (University of Washington) (40). The double mutants Δ*ldtJ* Δ*mrcA* and Δ*ldtJ* Δ*mrcB* were generated by a two-steps protocol based on Cre recombination followed by natural transformation of genomic DNA (40). Briefly, the tetracycline resistant cassette of the recipient single mutant was removed by site-specific recombination. Subsequently, purified genomic DNA of the donor mutant was mixed 1:1 with the recipient mutant grown to early logarithmic phase (OD_600_ 0.2 – 0.3), spotted on freshly prepared LB agar and incubated for 4 hours at 37°C. After incubation the bacteria were resuspended in PBS and spotted on LB agar supplemented with tetracycline for selection and incubated at 28°C overnight. Mutations were confirmed by PCR and verified by Whole Genome Sequencing (WGS). Two independent double mutant isolates were characterized and only one is shown. The *ldtJ* (ABUW_118) and *mrcA* (ABUW_0289) genes were cloned into the BamHI/XbaI restriction sites of the replicative plasmid pJMP3665^HygR^ (41). pJMP3665*::ldtJ* was introduced into *E. coli* DAP^-^mating strain, and an overnight culture was mixed at equal ratios with an overnight culture of the Δ*ldtJ* Δ*mrcA* recipient mutant for conjugative transfer. The cell mixture was spotted on an LB plate supplemented with DAP and incubated at 37°C for 3 hours. After this, grown bacteria were resuspended in PBS and spotted on LB agar supplemented with 250 µg/ml hygromycin for selection and incubated at 37°C overnight (41). LdtJ expression was induced by adding 1 mM of IPTG. pJMP3665*::mrcA* was introduced the Δ*ldtJ* Δ*mrcA* recipient mutant by electroporation of the ligation product. Briefly, the recipient culture was grown at 37°C until mid-logarithmic phase (OD_600_ ∼0.4). After this, cells were washed five times with 10% ice-cold glycerol. 500 ng of ligation DNA was electroporated into the recipient cells in a 2-mm cuvette at 1.8 kV. Cells were recovered in LB broth in shaking conditions for 1 hour at 37 °C and plated on an LB agar plate supplemented with 250 µg/ml hygromycin. PBP1A expression was induced by adding 10 µM IPTG.

### Construction of ATCC 17978 Δ*ldtJ* Δ*mrcA* double mutant and ectopic expression of *ldtJ* and *mrcA*

ATCC 17978 Δ*ldtJ* Δ*mrcA* double mutant was generated as previously described (17,29,42,43) by introducing the recombinase plasmid pMMB67EH^TetR^ encoding REC*_Ab_* in the Δ*mrcA* single mutant. Expression of the REC*_Ab_* was induced by adding 2 mM IPTG and cells were grown at 37°C until mid-logarithmic phase (OD_600_ ∼0.4-0.5). After washing with 10% ice-cold glycerol, the linear PCR product targeting *ldtJ* with 125 bp region of homology and containing the kanamycin resistance cassette FLP recombination target (FRT) was electroporated into this mutant. Cells were recovered in LB broth in shaking conditions with 2 mM IPTG for 4 hours and plated on LB agar supplemented with 25 µg/ml kanamycin for selection. The mutation was verified by PCR. Mutants were then treated to remove the pMMB67EH^TetR^::REC*_Ab_* by restreaking on LB agar containing 2 mM NiCl_2_ (29,42,43). Mutants that lost the tetracycline resistance cassette were then electroporated with pMMB67EH^TetR^ carrying the FLP recombinase to excide the kanamycin cassette after recovering for 1 hour in LB broth and plating on LB agar with 10 µg/ml tetracycline and 2 mM IPTG to induce the expression of FLP. The kanamycin cassette excision was verified by PCR.

To ectopically express the *ldtJ* (A1S_2371) and *mrcA* (A1S_3196-97) genes, pMMB*::ldtJ* and pABBR*::mrcA* (with its native promoter) were electroporated in the of ATCC 17978 Δ*ldtJ*Δ*mrcA* double mutant to generate Δ*ldtJ*Δ*mrcA*/pLdtJ and Δ*ldtJ*Δ*mrcA*/pPBP1A strains, respectively. The Δ*ldtJ*Δ*mrcA*/pLdtJ strain was selected with 25 µg/ml kanamycin and *ldtJ* expression induced with 2 mM IPTG whereas the Δ*ldtJ*Δ*mrcA*/pPBP1A strain was selected with 10 µg/ml tetracycline and *mrcA* induction was constitutive.

### Growth curves and viability

Overnight cultures of AB5075 were diluted 1:50 in 12.5 ml of LB broth and pre-cultured to early logarithmic phase (OD_600_ 0.2-0.3) at 37°C with continuous shaking. At this point, pre-cultures were back-diluted 1:50 in 50 ml of LB broth, incubated at 37°C with continuous shaking and the growth was monitored by manually measuring the optical density at 600 nm every 30 minutes for 6 hours. Samples for viable counts were collected every 1.5 hours, serially diluted and plated on LB agar. Plates were incubated overnight at 37°C and the colony forming units (CFUs) were counted. Growth and viability were plotted using GraphPad Prism 10 (version 10.2.3). Each experiment was independently performed three times, and representative datasets are shown.

Overnight cultures of strain ATCC 17978 were back-diluted to an initial OD_600_ of 0.01 and inoculated into 5 ml of LB broth in triplicate. Cultures were incubated at 37°C with continuous shaking. Optical density at 600 nm was manually measured every hour for 10 hours. Growth curves were generated using GraphPad Prism 10 (version 10.2.3). Each experiment was independently performed three times, and representative datasets are shown.

### Microscopy and image analysis

Overnight cultures were diluted to an OD_600_ of 0.05 and grown in LB broth in shaking conditions at 37°C until reaching an OD_600_ of ∼0.4–0.5 (mid-logarithmic phase). The appropriate antibiotic and IPTG were added to the grown strains. Cells were harvested and washed in LB before being incubated with 10 mM HADA (7-hydroxycoumarincarbonylamino-D-alanine, Thermo Fisher Scientific) in LB broth for 20 min at 37°C, with shaking in dark conditions, to fluorescently label the sites of active peptidoglycan biosynthesis. After further washes, cells were resuspended in 1X phosphate-buffered saline (PBS) and fixed with 16% paraformaldehyde (PFA) and mounted on 1.5% agarose in 1X PBS pads on microscopy slides. Imaging was performed using an inverted Nikon Eclipse Ti-2 widefield epifluorescence microscope equipped with a Photometrics Prime 95B camera and a Plan Apo 100×, 1.45 NA objective lens. Images were acquired using NIS Elements software, and image analysis was carried out using the MicrobeJ plugin in Fiji. Cell length, width, aspect ratio, and fluorescence intensity were measured for at least 100 analyzed cells per experiment and plotted in GraphPad Prism 10 (version 10.2.3).

For FM 5-95 staining, cell cultures were collected at mid-logarithmic phase (OD_600_ ∼0.4), harvested by centrifugation and resuspended in 100 µl of 5% PFA solution (Immunofix, Bio-Optica). To stain and visualize membranes, cells were incubated with the FM 5-95 dye at a final concentration of 5 µg/ml for 10 minutes at room temperature, washed twice in PBS and resuspended in a 1:4 ratio of ProLong^TM^ Gold antifading (Thermo Fisher Scientific) and visualized at the microscope. Imaging was performed using an inverted Nikon Eclipse Ti2 equipped with a CREST V3 X-light Spinning Disc confocal module, Lumencor CELESTA multi-line lasers, a monochromatic sCMOS camera, and a monochromatic EMCCD camera for widefield and spinning disc acquisitions. Images were captured with an exposure time of 80 ms and an emission wavelength of 683 nm. For the widefield fluorescence Spinning Disk confocal channel modality, imaging parameters included an excitation wavelength of 488 nm selected using the Full Multiband Penta filter (DAPI/FITC/TRITC/Cy5/Cy7: 391/477/549/639/741), a laser intensity of 25%, and an exposure time of 150 ms. This configuration resulted in an emission wavelength of 684 nm collected using the Cy5 filter (685/40). Images were acquired using NIS Elements AR 5.42 software.

### Genomic extraction and whole genome sequencing analysis

Genomic DNA from AB5075 was isolated using a EasyDNA genomic kit (Invitrogen) and subject to Illumina Whole Genome Sequencing (SeqCenter, Pittsburgh). Mean genome coverage was 103.1-fold, (SD 17.8). Gene inactivations by transposon and background and potential suppressor mutations were identified by use of the breseq pipeline (44), including Bowtie 2 for sequence alignment (45), using the default settings. Because the source of the mutant library was the ABUW_5075 transposon resource (40), genome file <u>CP008706</u> of *A. baumannii* strain AB5075-UW was used for sequence read alignment. Briefly, high-confidence mutations, flagged as sequence reads differing from the reference alignment and missing coverage and new junctions were assigned priority, if not present in the sequence of the parent strain, although marginal calls or missing coverage were evaluated both visually and in the fastq files when present at high frequency in the same gene or enzymatic pathway, when the same gene was mutated in more than one independent clone, and specifically when the base change or changes would result in a drastic change in the protein i.e., stop codons or change of reading frame were followed up, while intergenic changes, synonymous codon changes and changes that would lead to similar amino acid incorporation were not.

### Colony spot susceptibility assays

Overnight cultures were serially diluted from 10⁰ to 10⁻^5^ and spotted onto LB agar plates containing varying concentrations of detergent (SDS or SDS/EDTA) or antibiotic (rifampin, colistin or vancomycin). Plates were incubated at 37°C and growth was monitored to assess strain susceptibility under each condition.

### Peptidoglycan isolation and muropeptide analysis

AB5075 cultures were grown in 400 ml of LB medium until they reached either mid-logarithmic or stationary phase. Cells were harvested at 4°C and resuspended in 6 ml of pre-chilled water. The suspensions were dropped into 6 ml of boiling 8% SDS solution and boiled for 30 minutes. Muropeptides were isolated using a previously established method (8). In short, peptidoglycan was digested with the muramidase Cellosyl (Hoechst, Frankfurt am Main, Germany) to release muropeptides, which were then reduced with sodium borohydride. Separation of the reduced muropeptides was performed on a reversed-phase Prontosil 120-3-C18 AQ column (250 × 4.6 mm, 3 µm; Bischoff, Leonberg, Germany). The eluates were monitored at 205 nm by UV absorbance. Peaks were identified by comparison with previously published chromatographic profiles of *A. baumannii* 17978 (17,22,29), and novel peaks were further identified through tandem mass spectrometry (MS/MS). For each condition, mean values and variability were determined from two independent biological replicates.

### Generation of PBP1A-FLAG and LdtJ-FLAG fusion constructs

The primers used to generate PBP1A-FLAG and LdtJ-FLAG are listed in Table S3. The *mrcA* gene encoding PBP1A and the *ldtJ* gene encoding LdtJ from *A. baumannii* 17978 were amplified, purified, and cloned into the KpnI/BamHI sites of pMMB^KanR^ (17) to create pPBP1A-FLAG and pLdtJ-FLAG, respectively. Plasmids were then transformed into wild type cells and the respective single mutants, and the gene expression was induced with 1mM IPTG for co-immunoprecipitation.

### Co-immunoprecipitation and Western blotting

As previously described (46), overnight cultures were diluted to an initial optical density of OD_600_ 0.05 in 400 ml of LB broth and grown to late-logarithmic phase (OD_600_ ∼0.9) at 37°C. Cells from 100 ml culture were harvested, washed with 1X PBS, and incubated with 25 mM dithiobis-succinimidyl propionate (DSP, Thermo Fisher Scientific) cross-linker in 1 ml PBS for 1 hour in shaking conditions at 37°C. The reaction was quenched by incubation at room temperature with 400 μl of 1 M glycine with rocking for 15 minutes. Cells were then lysed and solubilized overnight at 4°C with rocking. Lysates were centrifuged at 10000 x g for 20 minutes and the supernatant was centrifuged twice to completely remove the pellet. 30 μl of anti-FLAG M2 affinity resin (Sigma) was added to the supernatant and incubated overnight at 4°C with rocking. The resin was then collected by centrifugation at 6000 x g for 15 minutes and washed 4 times with radioimmunoprecipitation assay (RIPA) buffer. Bound proteins were eluted by resuspending the resin in 100 μl SDS-PAGE loading buffer supplemented with 5% β-mercaptoethanol (BME), followed by boiling at 95°C for 7 minutes and samples were used for Western blotting.

Western blotting was performed by transferring proteins onto polyvinylidene fluoride (PVDF) membranes (Thermo Fisher Scientific). Membranes were blocked with 5% milk for 2 hours, followed by incubation with primary antibodies ⍺-FLAG (1:1000), ⍺- PBP1A (1:1000), ⍺- LdtJ (1:500), ⍺- 6×His (1:500), and ⍺- RpoA (1:1000). Detection was carried out using horseradish peroxidase (HRP)-conjugated secondary antibodies: anti-rat HRP (1:10000; Thermo Fisher Scientific) for anti-FLAG, and anti-rabbit HRP (1:10000; Thermo Fisher Scientific) for all other primary antibodies.

### Isolation of LOS⁻ *A. baumannii* and determination of mutation frequency

Isolation of LOS⁻ *A. baumannii* colonies was performed as described previously (17,22). Overnight cultures at OD_600_ of 1.0 (∼10⁹ CFU) were harvested by centrifugation at 8,000 rpm for 2 min, washed once with 1 ml of LB broth, and plated on LB agar supplemented with colistin (10 µg/ml). After overnight incubation, isolated colistin-resistant colonies were repatched onto LB agar containing vancomycin (10 µg/ml), colistin (10 µg/ml), and no antibiotic as control. Colonies resistant to colistin but sensitive to vancomycin were considered LOS-. The LOS- recovery frequency was calculated as the ratio of LOS- colonies to the colony forming units count in 100 μl.

### LOS staining and analysis

LOS analysis was done as previously reported (42). Overnight cultures were diluted to an OD_600_ of 0.1 in 5 ml LB broth and grown at 37°C with shaking until reaching an OD_600_ of ∼1. 1 ml of cells were harvested and resuspended in 100 µl 1X Sample Buffer (prepared from 4X LDS Sample Buffer, 4% β-mercaptoethanol, and water) and boiled for 10 minutes before incubating at 55°C overnight with proteinase K. The following day, samples were boiled for 10 minutes and subjected to SDS-PAGE. Gels were fixed and stained following the Pro-Q Emerald 300 Lipopolysaccharide Gel Stain Kit (Thermo Fisher Scientific, P20495).

### Determination of minimal inhibitory concentrations (MICs)

MIC determination was done as previously described (47), using E-test strips (Liofilchem). The Δ*ldtJ* Δ*mrcA* colistin-resistant LOS- overnight culture was concentrated 10X and spread uniformly on LB agar plates using a sterile cotton swab, completely covering of the agar surface. Once the plates dried, an E-test strip was placed onto the surface in sterile conditions. Plates were incubated at 37°C for 18–20 hours.

## Supporting information

Furlan et al_PBP1a LdtJ manuscript_full text, tables, figures

## Acknowledgments

We thank Alessandra Polissi and Alessandra Martorana for stimulating ideas and discussion. We would also like to thank Jason Peters and his laboratory for sharing the hygromycin replicative plasmids for complementation in AB5075. We thank Philip Rather and his laboratory for generously sharing their *Acinetobacter baumannii* AB5075 mutagenesis and natural transformation protocols, which facilitated construction of the mutants used in this study.

Work in the O.M. lab was supported by the internal funding from the Interdisciplinary Center for Medical Sciences (CISMed) to O.M. Work in the J.M.B. lab was supported by the National Institutes of Health grants R01AI168159 and R35GM143053. Work in the W.V. lab was supported by the NIH grant R01AI168159. The work of B.F. was also supported by the FEMS Research and Training Grant.

F., S.N., H.G., M.B.W., W.V., J.M.B, and O.M. participated in the conception or design of the study; B.F., S.N., H.G., M.B.W., N.M., R.J.O.O, D.V., W.V., J.M.B. and O.M. participated in the acquisition, analysis, or interpretation of the data; B.F., S.N. and R.J.O.O. wrote the manuscript; and W.V., J.M.B. and O.M. made comments on and revisions to the manuscript.

## Supplementary Figures

**Figure S1.**
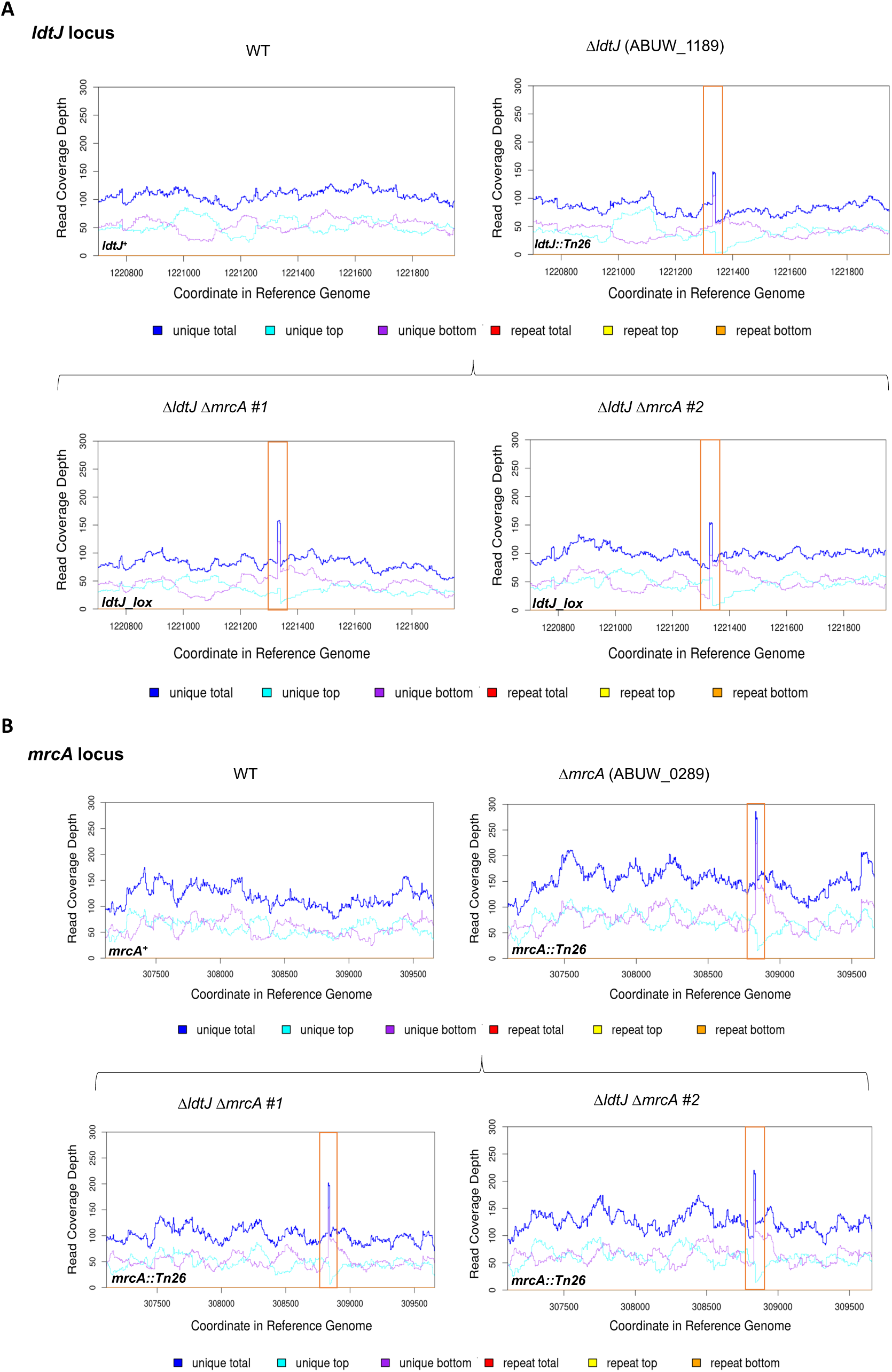
Whole-genome sequencing of *A. baumannii* AB5075 WT, Δ*ldtJ,* Δ*mrcA* and Δ*ldtJ* Δ*mrcA* double mutants. Whole-genome sequencing was performed to confirm the simultaneous inactivation of the *ldtJ* and *mrcA* genes and to verify the absence of suppressor mutations in the Δ*ldtJ* Δ*mrcA* double mutants. **(A)** Read coverage depth across the *ldtJ* locus. **(B)** Read coverage depth across the *mrcA* locus. Both *ΔldtJ ΔmrcA* #1 and *ldtJ ΔmrcA* #2 mutants showed an insertion (*tn* or *tnlox*) in the *ldtJ* locus and *mrcA* locus, confirming double inactivaction of *ldtJ* and *mrcA*.

**Figure S2.**
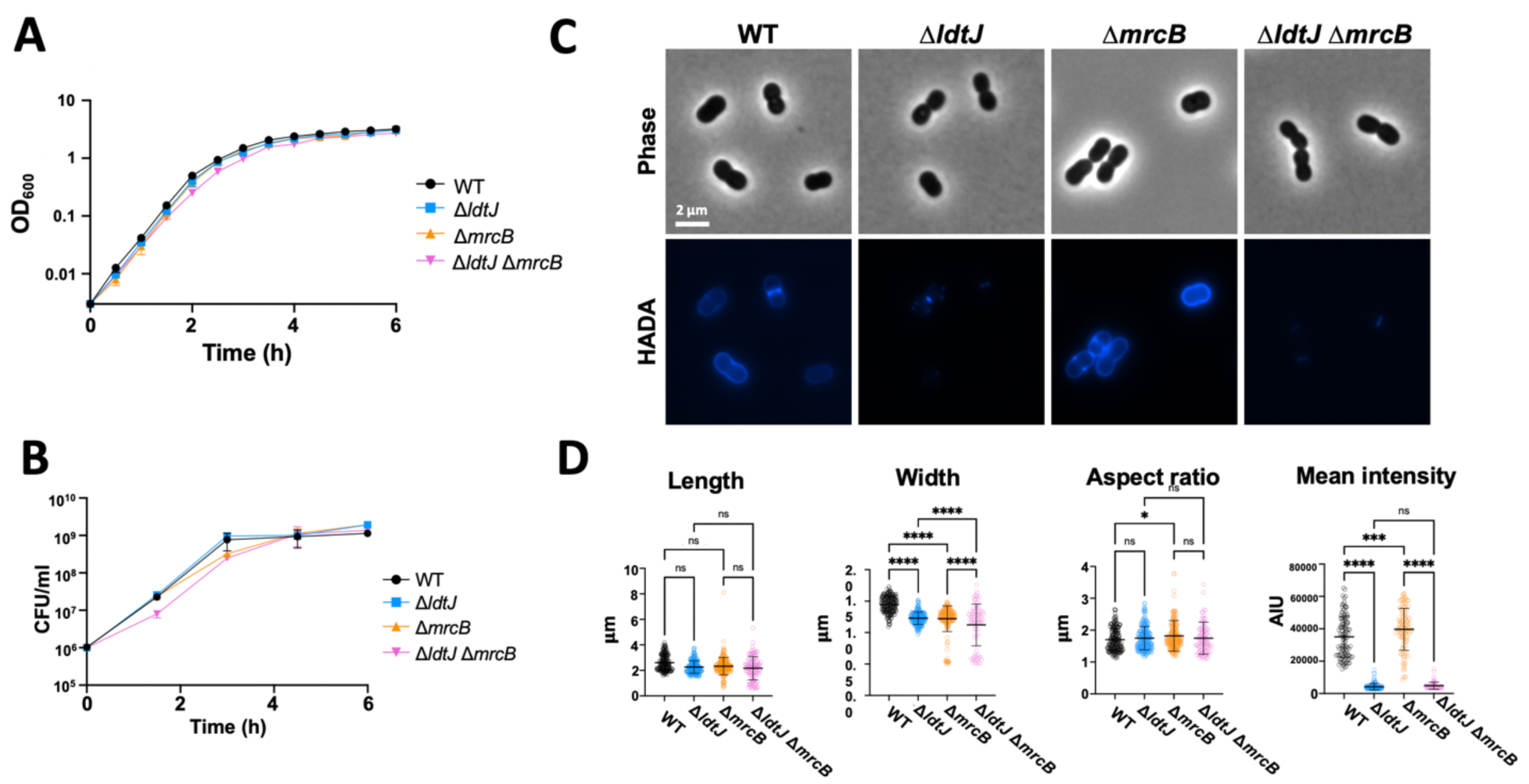
Characterization of the *A. baumannii* AB5075 Δ*ldtJ* Δ*mrcB* double mutant. **(A)** Growth (OD_600_) and **(B)** viability (CFU/ml) of wild type (WT), Δ*ldtJ*, Δ*mrcB,* and Δ*ldtJ* Δ*mrcB* mutants in LB. Each experiment was independently performed in triplicate. **(C)** Phase-contrast and D-amino acid (HADA) fluorescence microscopy of the WT and mutants in logarithmic phase of growth. **(D)** Quantification of cell length, width, aspect ratio, and HADA fluorescence intensity of each population (*n >*100 cells per strain), analyzed using ImageJ software. Statistical significance was determined using one-way ANOVA (\**P* < 0.05; \*\*\**P* < 0.001; \*\*\*\**P* < 0.0001; ns = not significant).

**Figure S3.**
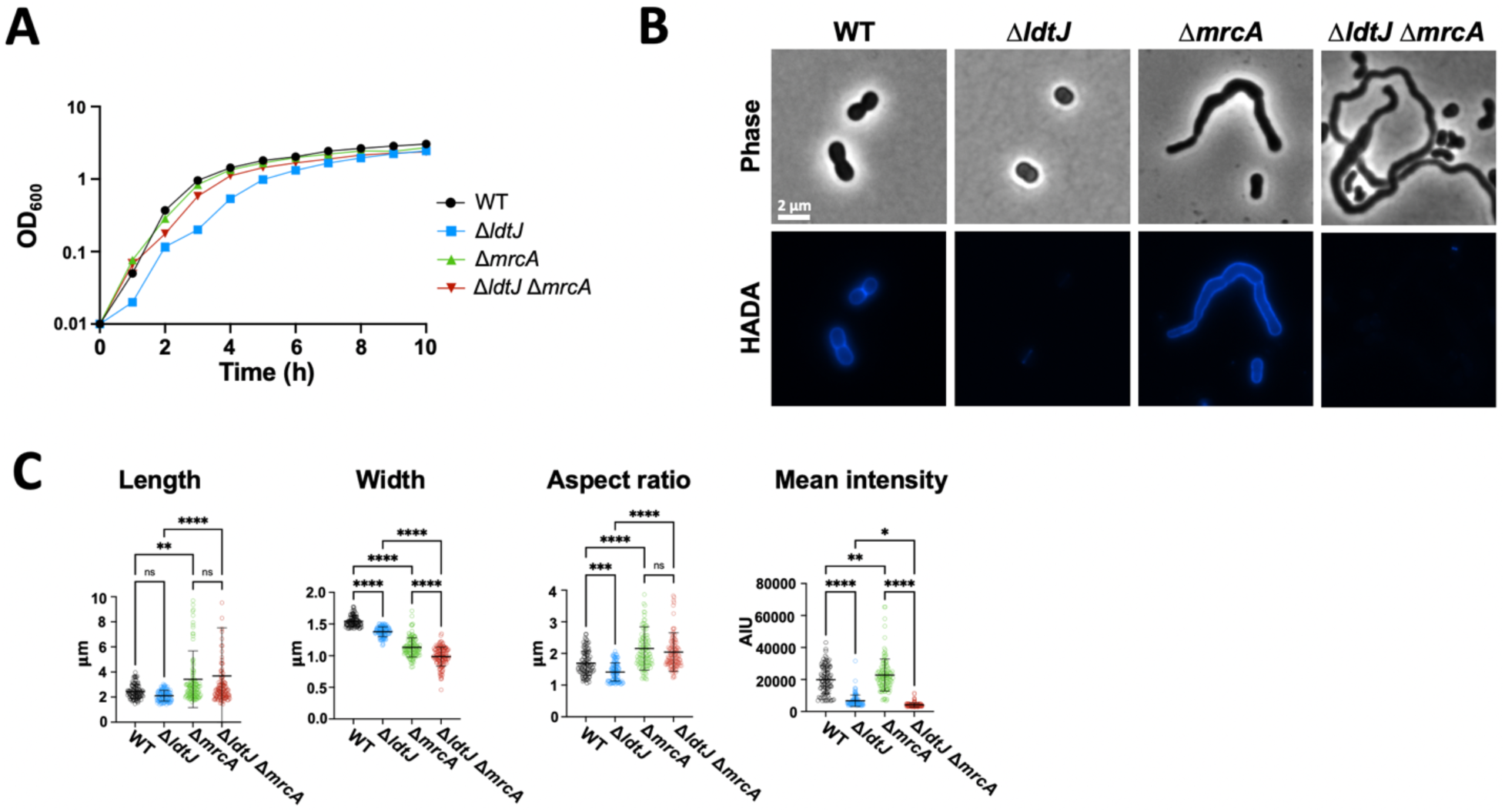
Characterization of the *A. baumannii* ATCC 17978 Δ*ldtJ* Δ*mrcA* double mutant. **(A)** Growth (OD_600_) of wild type (WT), Δ*ldtJ*, Δ*mrcA,* and Δ*ldtJ* Δ*mrcA* mutants in LB. Each experiment was independently performed in triplicate. **(B)** Phase-contrast and D-amino acid (HADA) fluorescence microscopy of the WT, Δ*ldtJ*, Δ*mrcA*, and Δ*ldtJ* Δ*mrcA* mutants. **(C)** Quantification of cell length, width, aspect ratio, and HADA fluorescence intensity of each population (*n >*100 cells per strain), analyzed using ImageJ software. Statistical significance was determined using one-way ANOVA (\**P* < 0.05; \*\**P* < 0.01; \*\*\**P* < 0.001; \*\*\*\**P* < 0.0001; ns = not significant).

**Figure S4.**
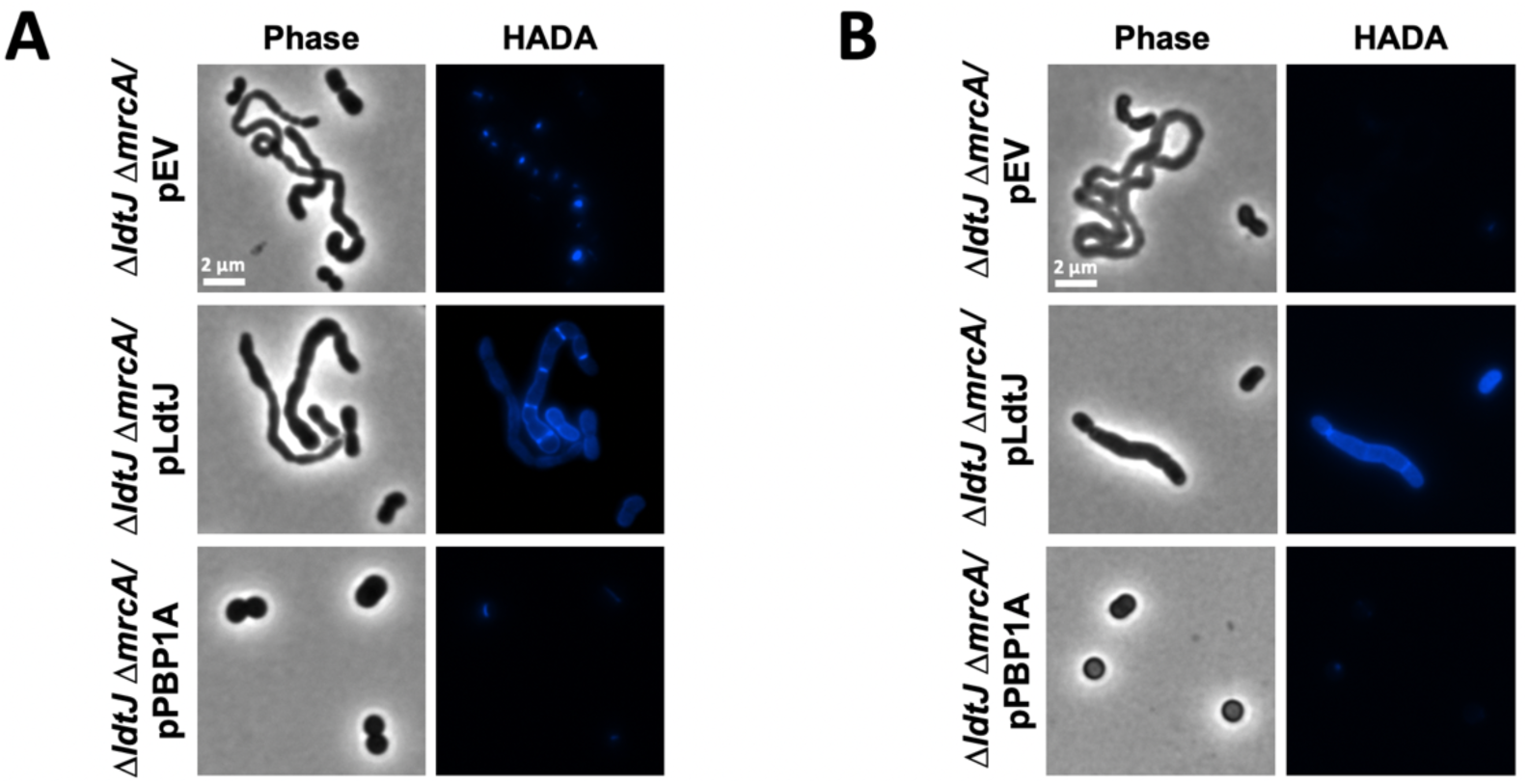
Ectopic production of PBP1A or LdtJ in the *A. baumannii* AB5075 and ATCC 17978 Δ*ldtJ* Δ*mrcA* double mutants restore the respective single mutants phenotype. **(A)** Phase-contrast and HADA fluorescence microscopy of the AB5075 Δ*ldtJ* Δ*mrcA* double mutant transformed either with the empty vector (pEV) or the plasmids expressing *mrcA* (pPBP1A) or *ldtJ* (pLdtJ) in logarithmic phase of growth. **(B)** Phase-contrast and HADA fluorescence microscopy of the ATCC 17978 Δ*ldtJ* Δ*mrcA* double mutant transformed either with the empty vector (pEV) or the plasmids expressing *mrcA* (pPBP1A) or *ldtJ* (pLdtJ) in logarithmic phase of growth.

**Figure S5.**
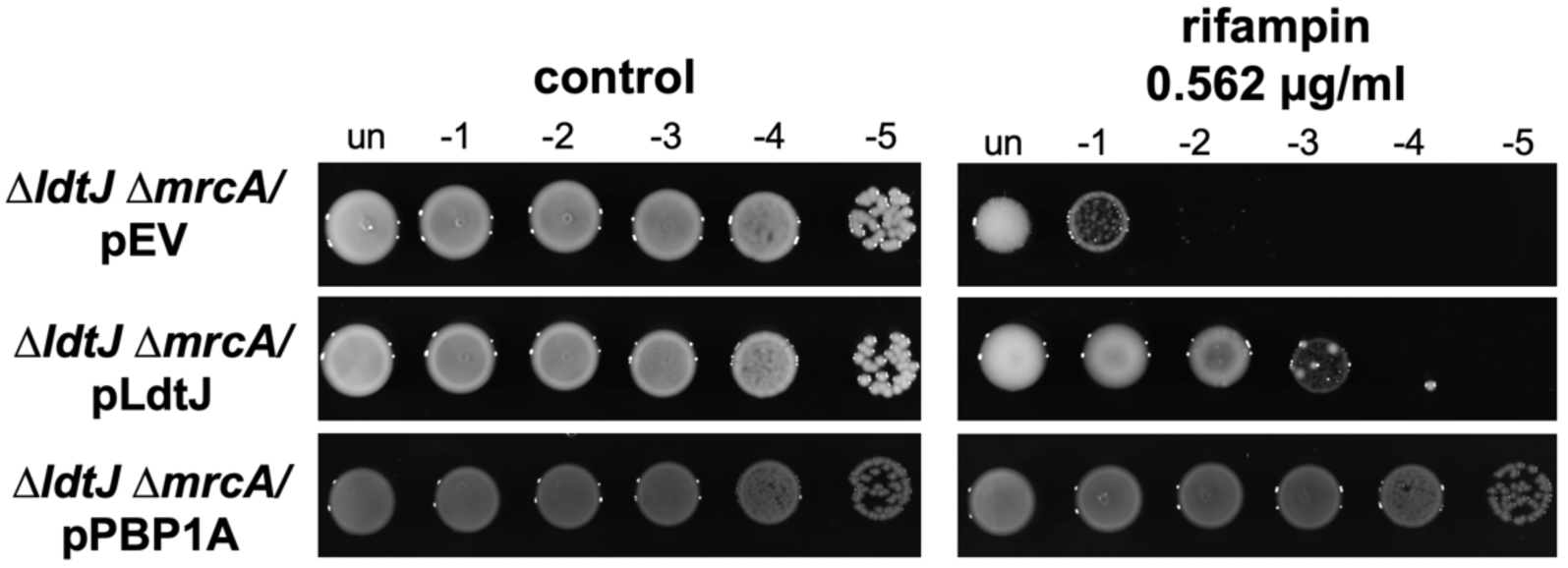
Ectopic production of PBP1a or LdtJ in the *A. baumannii* AB5075 Δ*ldtJ* Δ*mrcA* double mutant restore the ability to grow in the presence of rifampin. Colony spot assay of the Δ*ldtJ* Δ*mrcA* double mutant carrying pEV, pLdtJ or pPBP1A on LB agar with or without 0.562 µg/ml rifampin.

**Figure S6.**
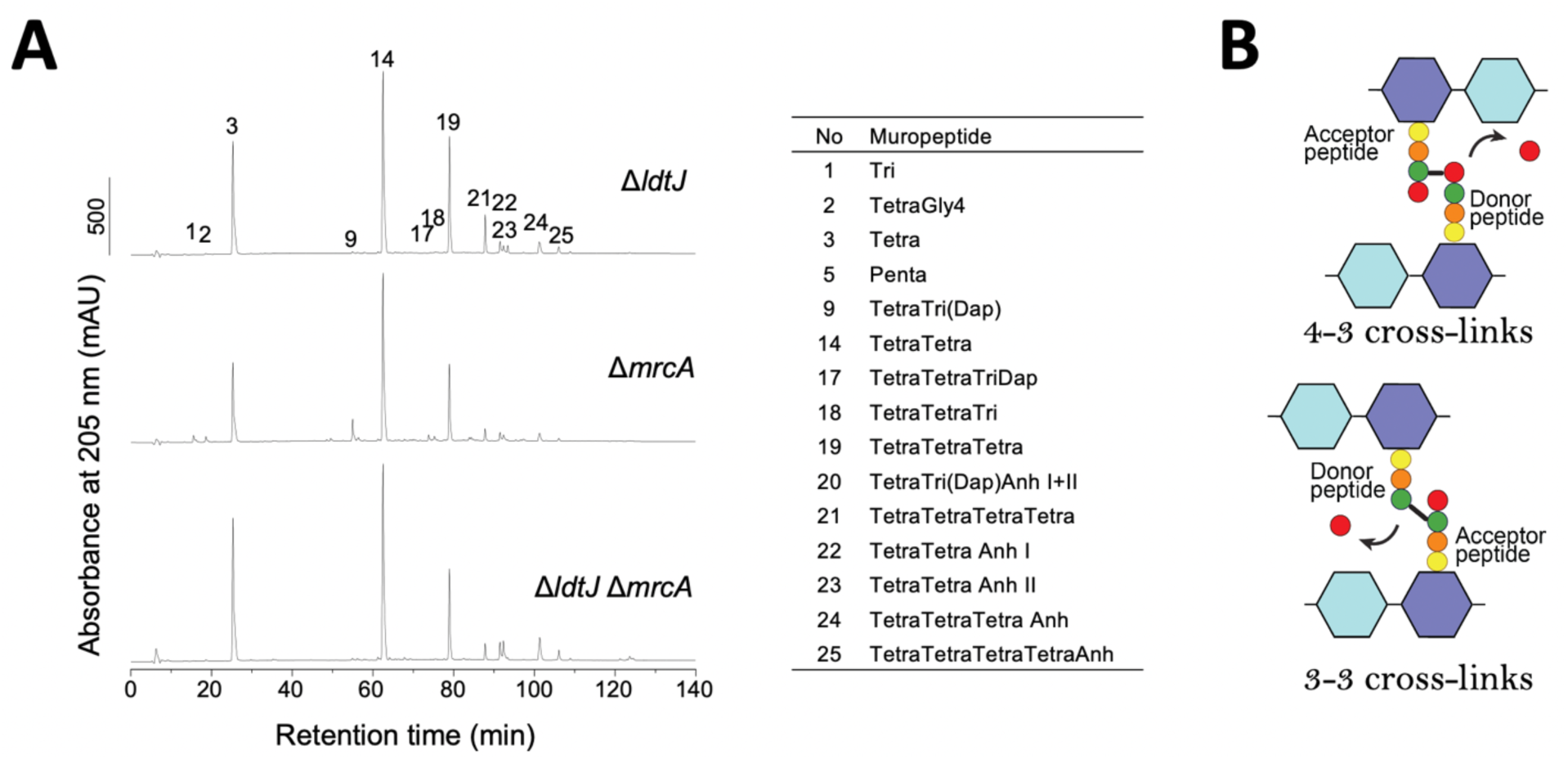
Cross-linkage changes are maintained in stationary phase. **(A)** PG isolated from *A. baumannii* AB5075 Δ*ldtJ*, Δ*mrcA* and the Δ*ldtJ* Δ*mrcA* mutants in stationary phases was analyzed by High Performance Liquid Chromatography (HPLC). **(B)** Schematic representation of 4-3 and 3-3 cross-links.

## Supplementary Tables

**Table S1:**
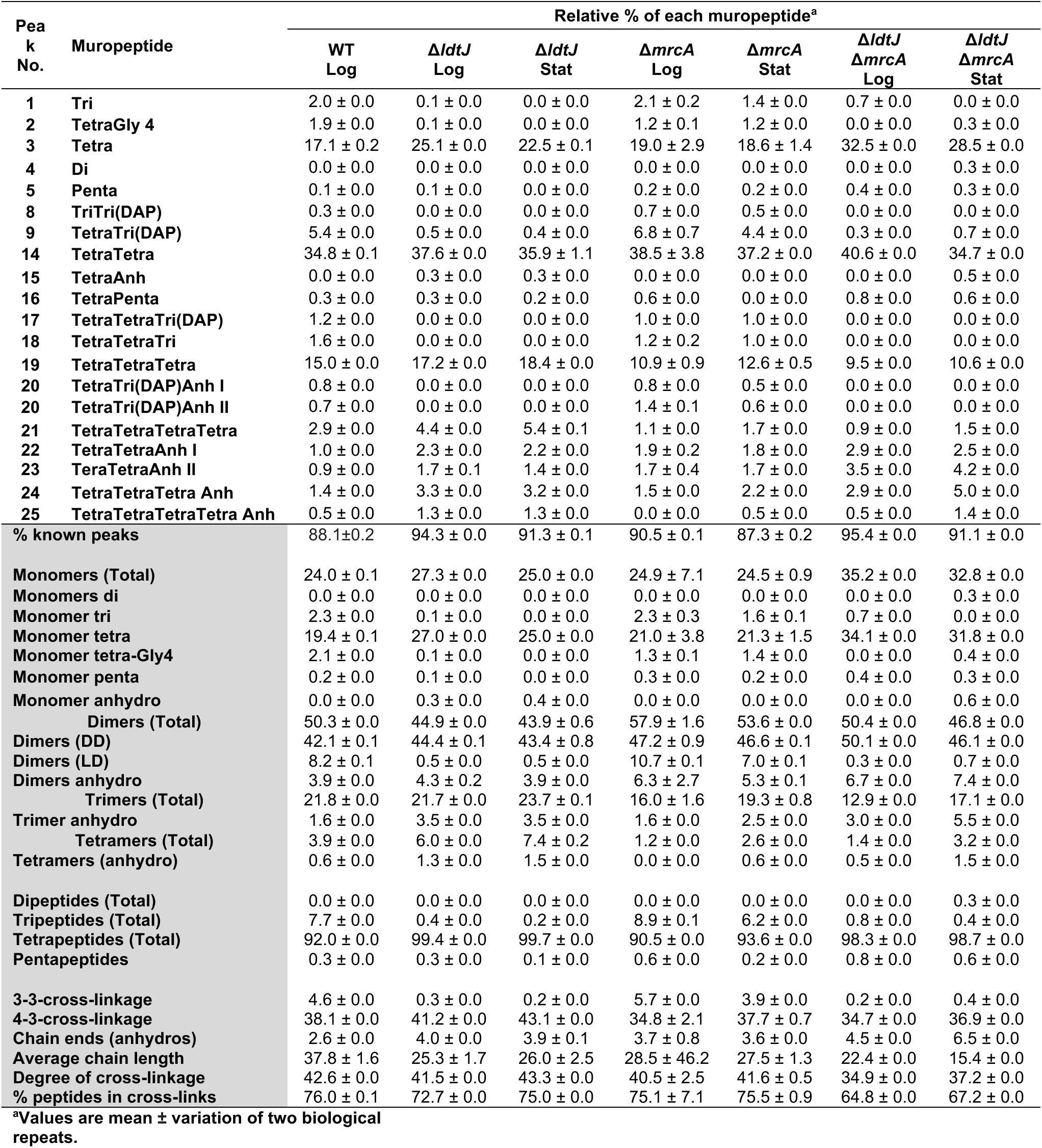
Muropeptide composition of wild type, Δ*ldtJ*, Δ*mrcA* and Δ*ldtJ* Δ*mrcA* in *A. baumannii* strain AB5075.

**Table S2:**
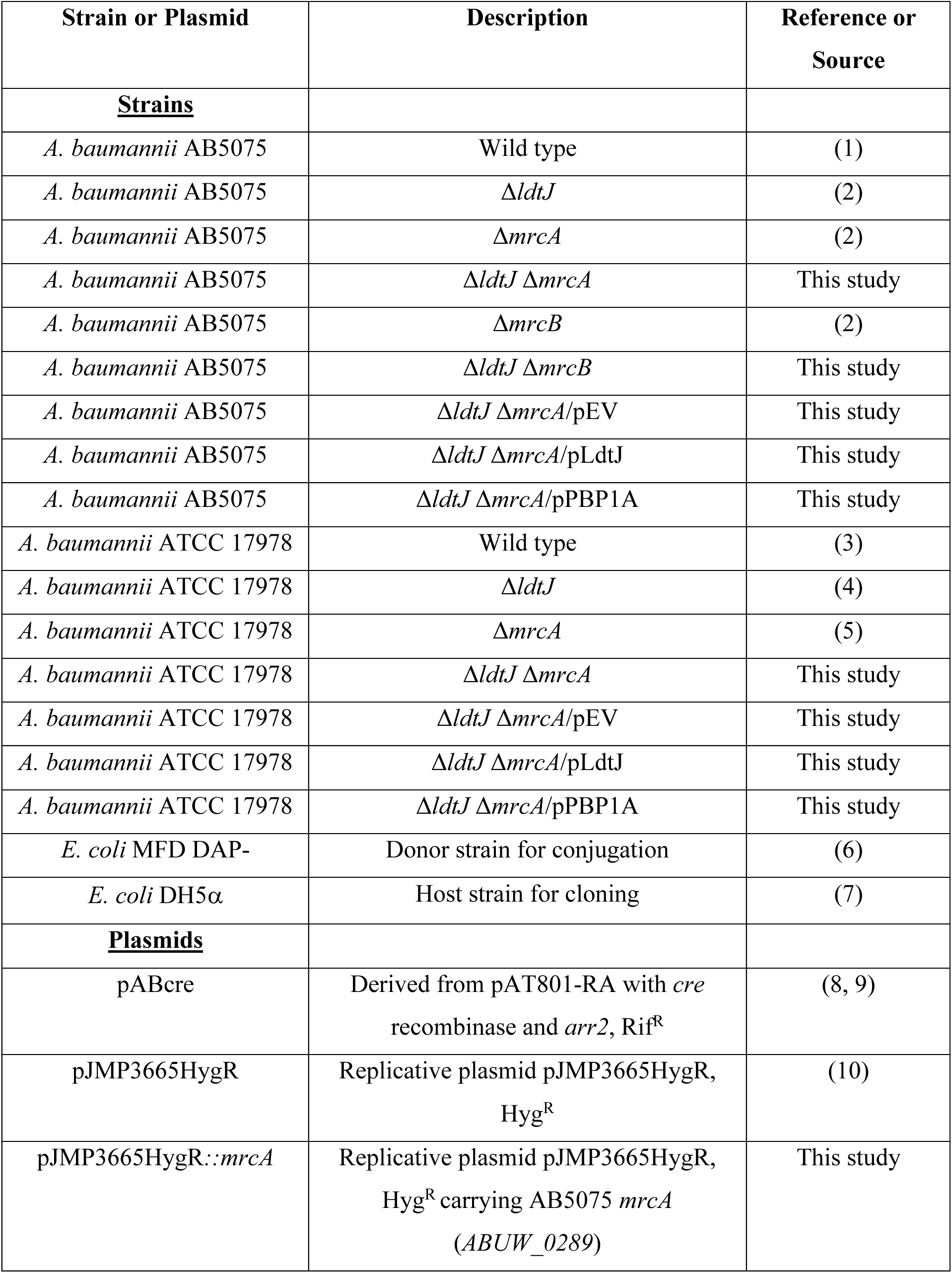
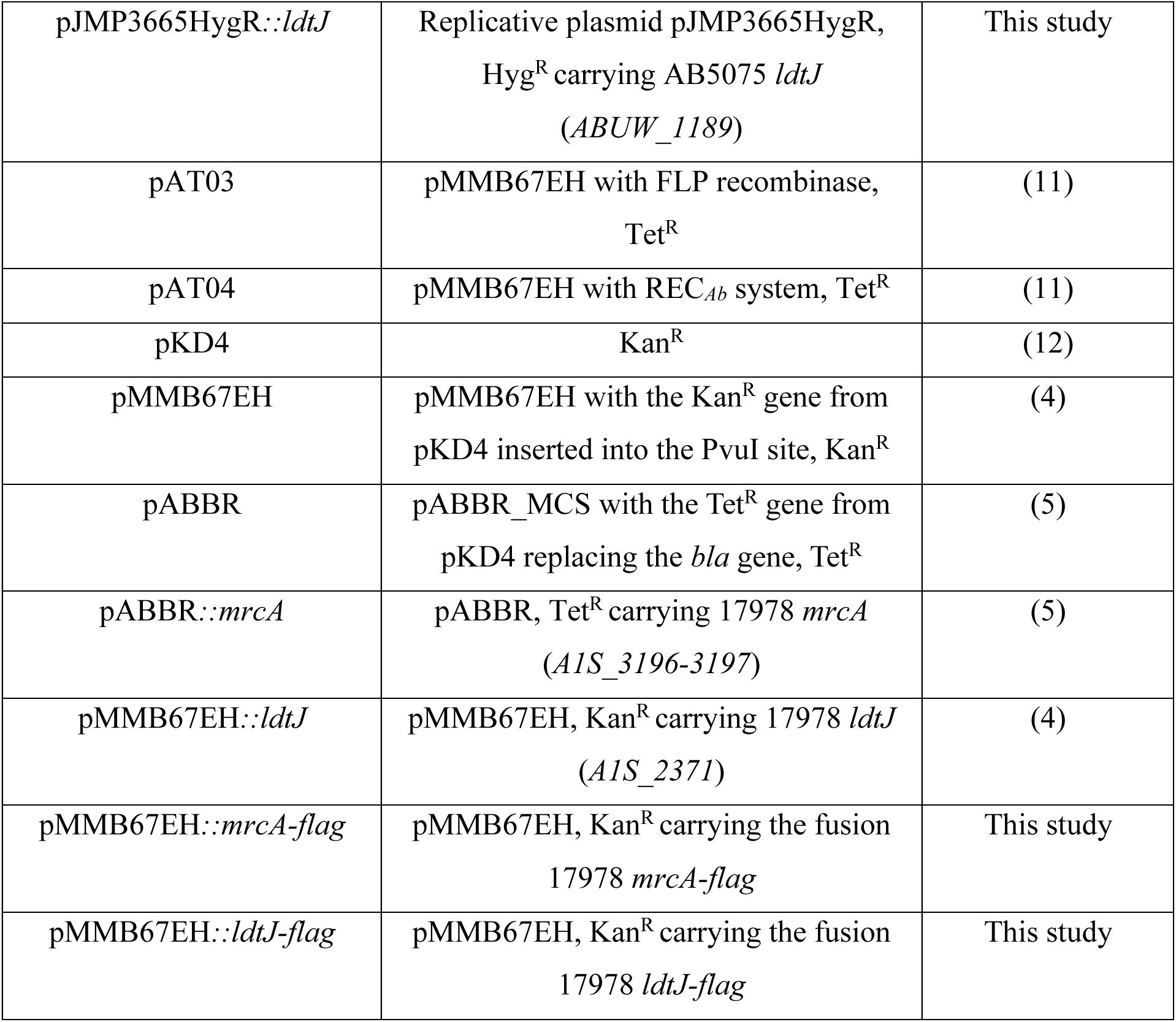
Strains and plasmids used in this study.

**Table S3:**
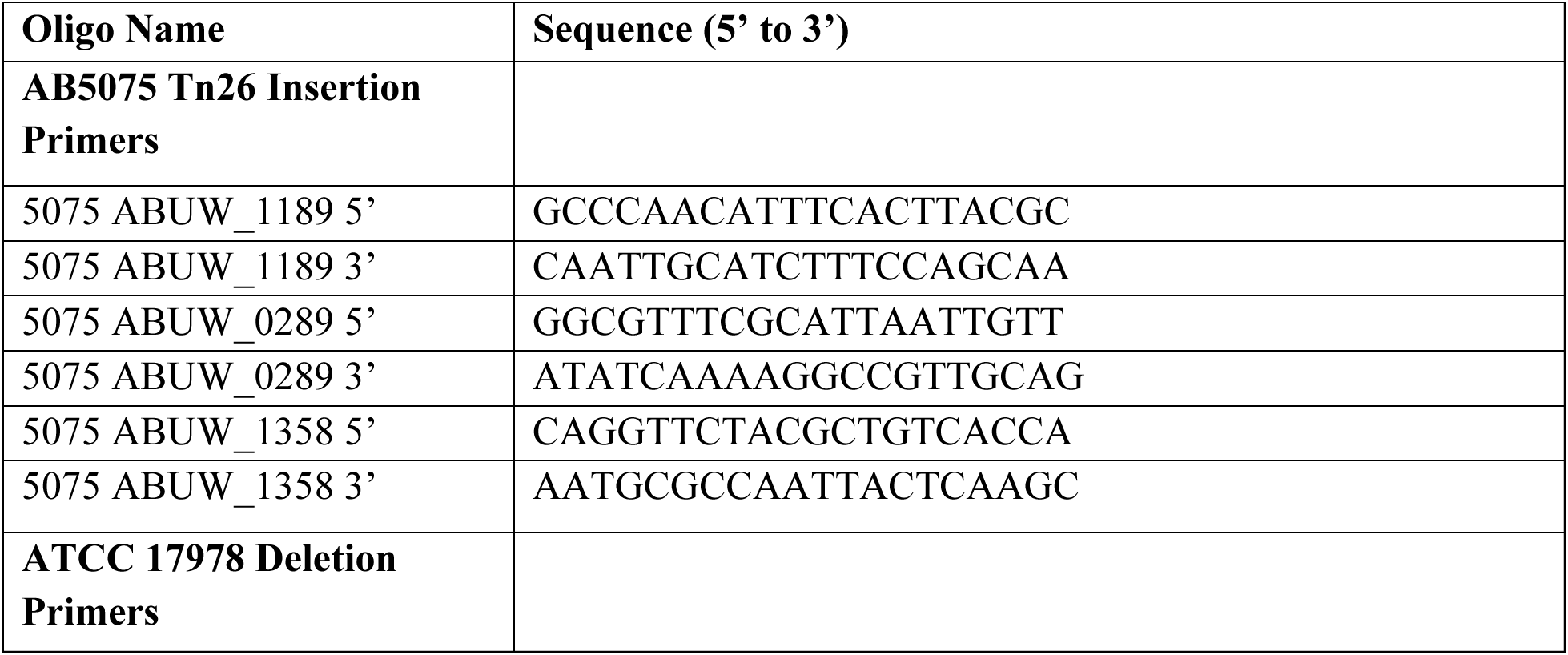
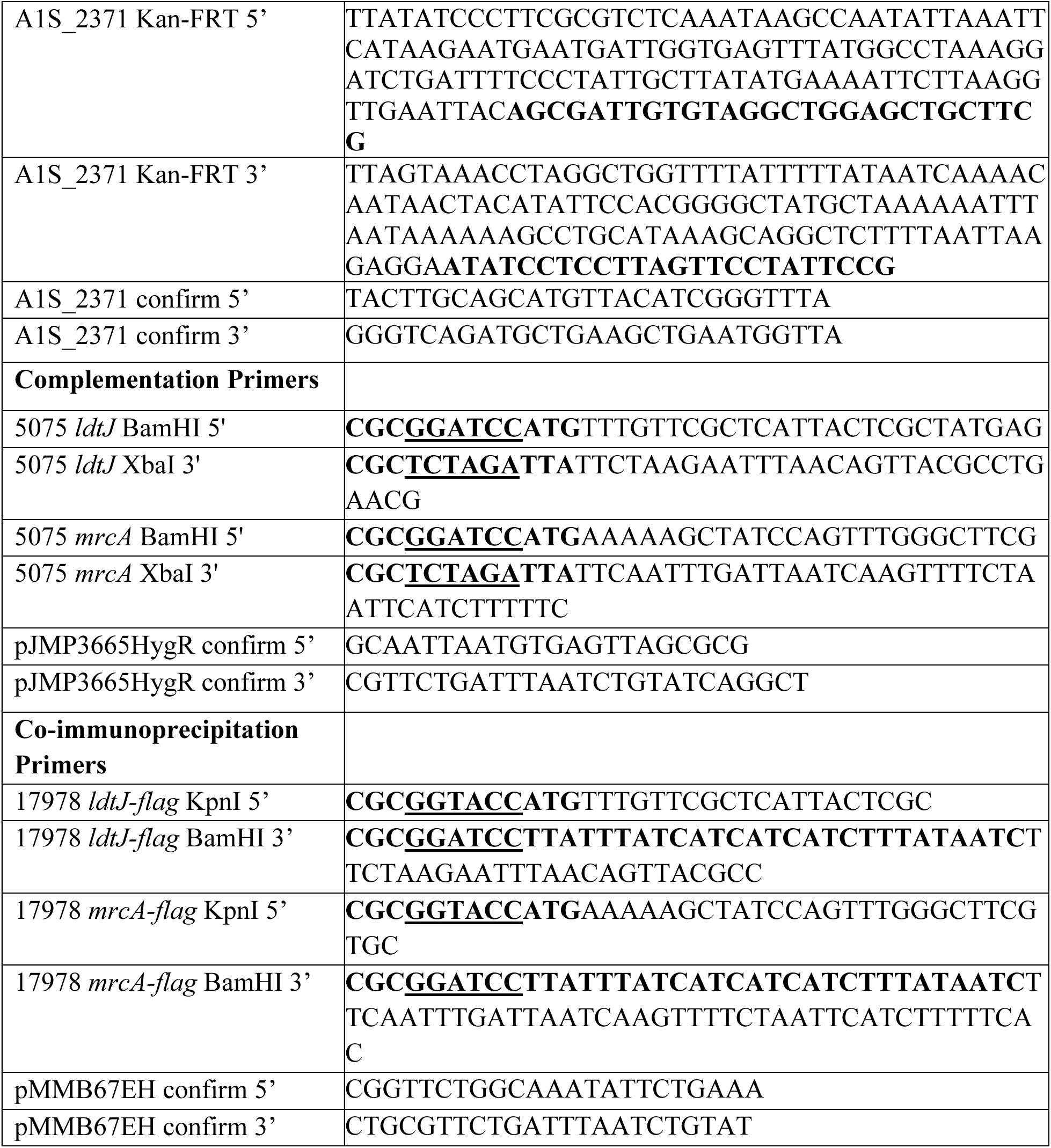
Primers used in this study.

