## Supplementary material for "PBP1A and LdtJ support cell envelope homeostasis and impact selection of Colistin-resistance in *Acinetobacter baumannii*": Furlan et al_PBP1a LdtJ manuscript_full text, tables, figures

*Co-correspondent authors

**Supplemental Figures and Figure' legends: FIGURES S1-S6**

**Supplemental Tables: TABLES S1-S4**

**Supplemental References**

### Supplementary Figures

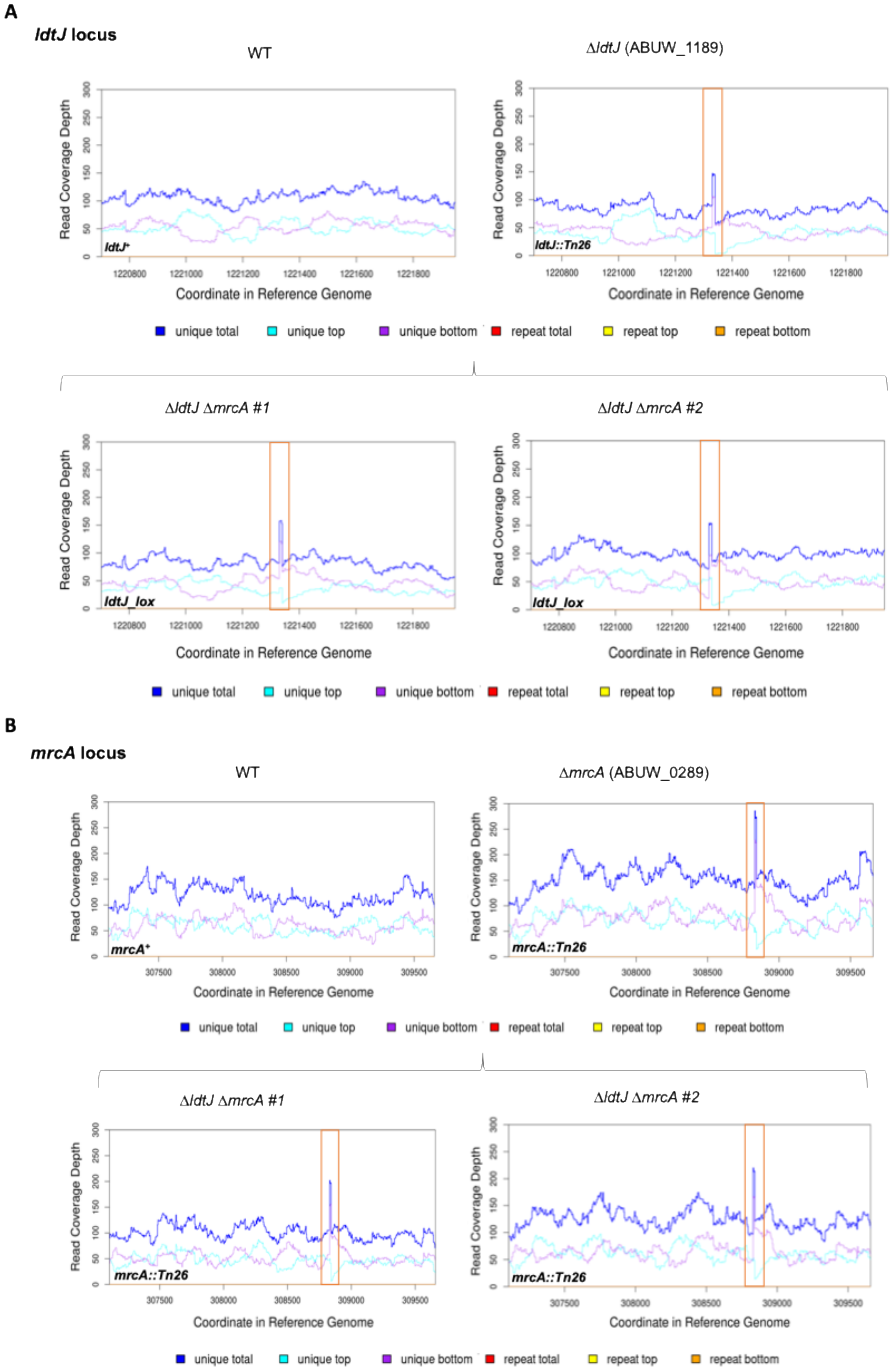

**Figure S1. Whole-genome sequencing of *A. baumannii* AB5075 WT, Δ*ldtJ,* Δ*mrcA* and Δ*ldtJ* Δ*mrcA* double mutants.** Whole-genome sequencing was performed to confirm the simultaneous inactivation of the *ldtJ* and *mrcA* genes and to verify the absence of suppressor mutations in the Δ*ldtJ* Δ*mrcA* double mutants. **(A)** Read coverage depth across the *ldtJ* locus. **(B)** Read coverage depth across the *mrcA* locus. Both Δ*ldtJ* Δ*mrcA* #1 and *ldtJ* Δ*mrcA* #2 mutants showed an insertion (*tn* or *tnlox*) in the *ldtJ* locus and *mrcA* locus, confirming double inactivation of *ldtJ* and *mrcA*.

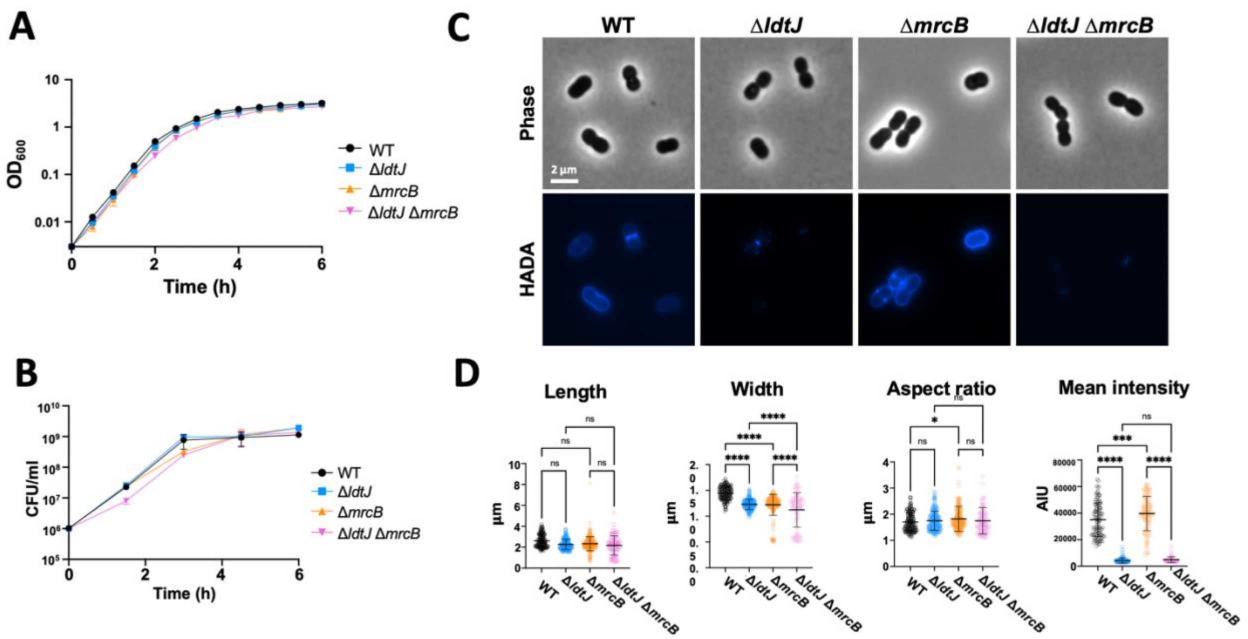


**Figure S2. Characterization of the *A. baumannii* AB5075 Δ*ldtJ* Δ*mrcB* double mutant. (A)** Growth ($OD_{600}$) and **(B)** viability (CFU/ml) of wild type (WT), Δ*ldtJ*, Δ*mrcB,* and Δ*ldtJ* Δ*mrcB* mutants in LB. Each experiment was independently performed in triplicate. **(C)** Phase-contrast and D-amino acid (HADA) fluorescence microscopy of the WT and mutants in logarithmic phase of growth. **(D)** Quantification of cell length, width, aspect ratio, and HADA fluorescence intensity of each population ($n$ >100 cells per strain), analyzed using ImageJ software. Statistical significance was determined using one-way ANOVA (*$P < 0.05$; ***$P < 0.001$; ****$P < 0.0001$; ns = not significant).

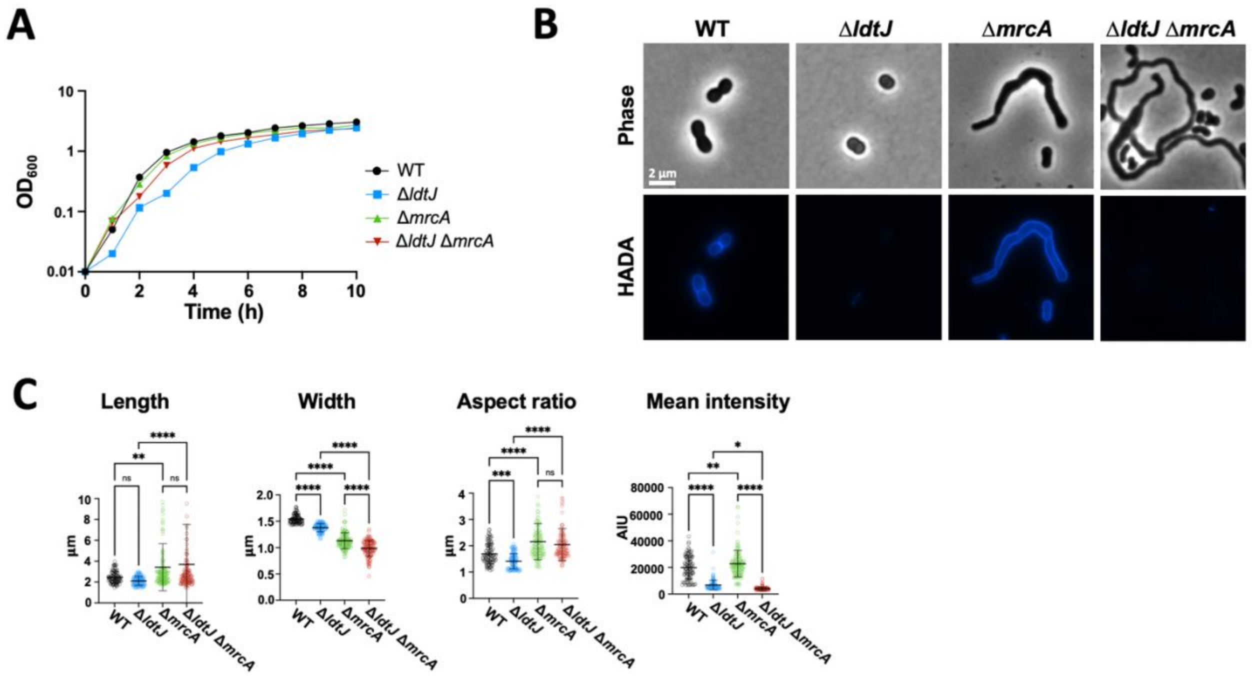


**Figure S3. Characterization of the *A. baumannii* ATCC 17978 Δ*ldtJ* Δ*mrcA* double mutant.** **(A)** Growth ($OD_{600}$) of wild type (WT), Δ*ldtJ*, Δ*mrcA,* and Δ*ldtJ* Δ*mrcA* mutants in LB. Each experiment was independently performed in triplicate. **(B)** Phase-contrast and D-amino acid (HADA) fluorescence microscopy of the WT, Δ*ldtJ*, Δ*mrcA*, and Δ*ldtJ* Δ*mrcA* mutants. **(C)** Quantification of cell length, width, aspect ratio, and HADA fluorescence intensity of each population (*n* >100 cells per strain), analyzed using ImageJ software. Statistical significance was determined using one-way ANOVA (**P* < 0.05; ***P* < 0.01; ****P* < 0.001; *****P* < 0.0001; ns = not significant).

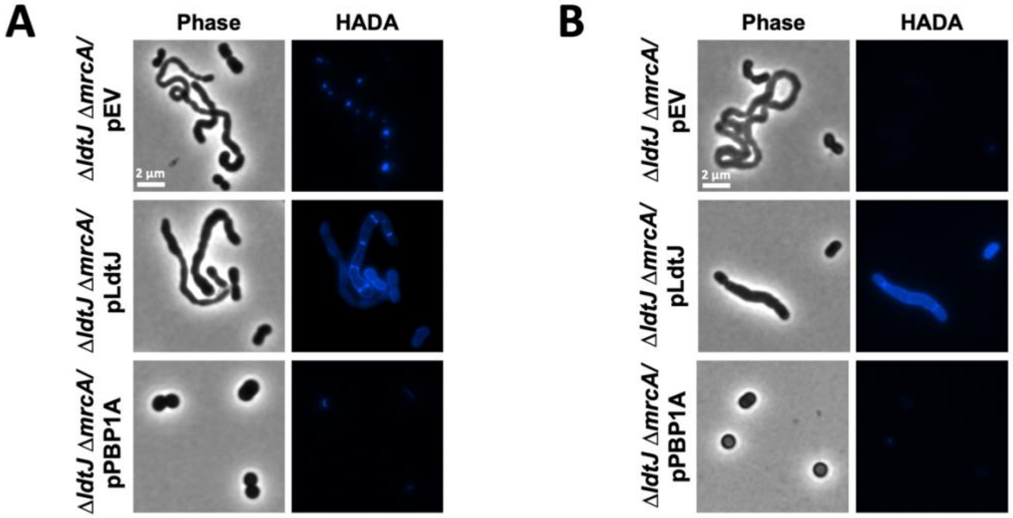


**Figure S4. Ectopic production of PBP1A or LdtJ in the *A. baumannii* AB5075 and ATCC 17978 Δ*ldtJ* Δ*mrcA* double mutants restore the respective single mutants phenotype. (A)** Phase-contrast and HADA fluorescence microscopy of the AB5075 Δ*ldtJ* Δ*mrcA* double mutant transformed either with the empty vector (pEV) or the plasmids expressing *mrcA* (pPBP1A) or *ldtJ* (pLdtJ) in logarithmic phase of growth. **(B)** Phase-contrast and HADA fluorescence microscopy of the ATCC 17978 Δ*ldtJ* Δ*mrcA* double mutant transformed either with the empty vector (pEV) or the plasmids expressing *mrcA* (pPBP1A) or *ldtJ* (pLdtJ) in logarithmic phase of growth.

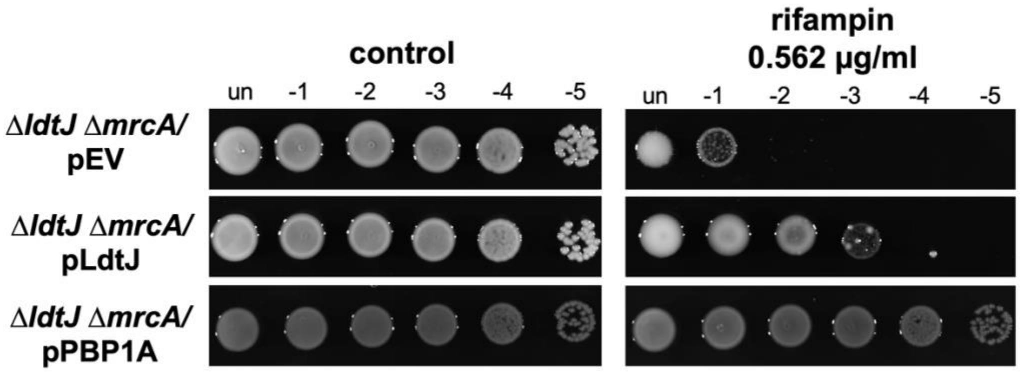


**Figure S5. Ectopic production of PBP1a or LdtJ in the *A. baumannii* AB5075 Δ*ldtJ* Δ*mrcA* double mutant restore the ability to grow in the presence of rifampin.** Colony spot assay of the Δ*ldtJ* Δ*mrcA* double mutant carrying pEV, pLdtJ or pPBP1A on LB agar with or without 0.562 µg/ml rifampin.

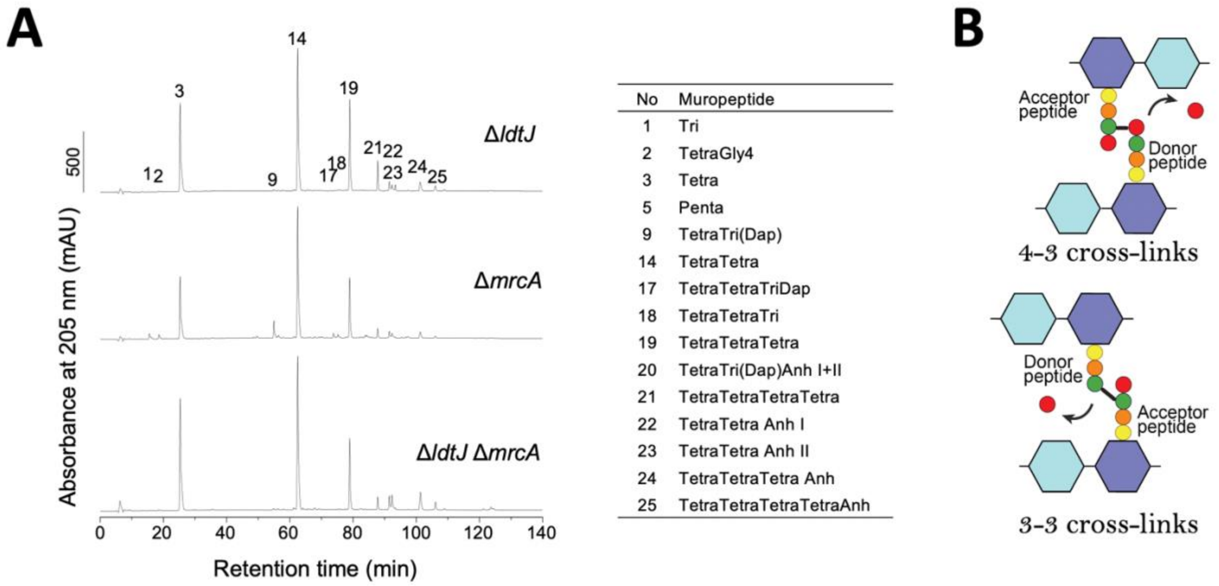


**Figure S6. Cross-linkage changes are maintained in stationary phase. (A)** PG isolated from *A. baumannii* AB5075 Δ*ldtJ*, Δ*mrcA* and the Δ*ldtJ* Δ*mrcA* mutants in stationary phases was analyzed by High Performance Liquid Chromatography (HPLC). **(B)** Schematic representation of 4-3 and 3-3 cross-links.

### Supplementary Tables

**Table S1:** Muropeptide composition of wild type, Δ*ldtJ*, Δ*mrcA* and Δ*ldtJ* Δ*mrcA* in *A. baumannii* strain AB5075.

| Peak No. | Muropeptide | Relative % of each muropeptide[a] | | | | | | |
|---|---|---|---|---|---|---|---|---|
| | | **WT Log** | **Δ*ldtJ* Log** | **Δ*ldtJ* Stat** | **Δ*mrcA* Log** | **Δ*mrcA* Stat** | **Δ*ldtJ* Δ*mrcA* Log** | **Δ*ldtJ* Δ*mrcA* Stat** |
| 1 | Tri | 2.0 ± 0.0 | 0.1 ± 0.0 | 0.0 ± 0.0 | 2.1 ± 0.2 | 1.4 ± 0.0 | 0.7 ± 0.0 | 0.0 ± 0.0 |
| 2 | TetraGly 4 | 1.9 ± 0.0 | 0.1 ± 0.0 | 0.0 ± 0.0 | 1.2 ± 0.1 | 1.2 ± 0.0 | 0.0 ± 0.0 | 0.3 ± 0.0 |
| 3 | Tetra | 17.1 ± 0.2 | 25.1 ± 0.0 | 22.5 ± 0.1 | 19.0 ± 2.9 | 18.6 ± 1.4 | 32.5 ± 0.0 | 28.5 ± 0.0 |
| 4 | Di | 0.0 ± 0.0 | 0.0 ± 0.0 | 0.0 ± 0.0 | 0.0 ± 0.0 | 0.0 ± 0.0 | 0.0 ± 0.0 | 0.3 ± 0.0 |
| 5 | Penta | 0.1 ± 0.0 | 0.1 ± 0.0 | 0.0 ± 0.0 | 0.2 ± 0.0 | 0.2 ± 0.0 | 0.4 ± 0.0 | 0.3 ± 0.0 |
| 8 | TriTri(DAP) | 0.3 ± 0.0 | 0.0 ± 0.0 | 0.0 ± 0.0 | 0.7 ± 0.0 | 0.5 ± 0.0 | 0.0 ± 0.0 | 0.0 ± 0.0 |
| 9 | TetraTri(DAP) | 5.4 ± 0.0 | 0.5 ± 0.0 | 0.4 ± 0.0 | 6.8 ± 0.7 | 4.4 ± 0.0 | 0.3 ± 0.0 | 0.7 ± 0.0 |
| 14 | TetraTetra | 34.8 ± 0.1 | 37.6 ± 0.0 | 35.9 ± 1.1 | 38.5 ± 3.8 | 37.2 ± 0.0 | 40.6 ± 0.0 | 34.7 ± 0.0 |
| 15 | TetraAnh | 0.0 ± 0.0 | 0.3 ± 0.0 | 0.3 ± 0.0 | 0.0 ± 0.0 | 0.0 ± 0.0 | 0.0 ± 0.0 | 0.5 ± 0.0 |
| 16 | TetraPenta | 0.3 ± 0.0 | 0.3 ± 0.0 | 0.2 ± 0.0 | 0.6 ± 0.0 | 0.0 ± 0.0 | 0.8 ± 0.0 | 0.6 ± 0.0 |
| 17 | TetraTetraTri(DAP) | 1.2 ± 0.0 | 0.0 ± 0.0 | 0.0 ± 0.0 | 1.0 ± 0.0 | 1.0 ± 0.0 | 0.0 ± 0.0 | 0.0 ± 0.0 |
| 18 | TetraTetraTri | 1.6 ± 0.0 | 0.0 ± 0.0 | 0.0 ± 0.0 | 1.2 ± 0.2 | 1.0 ± 0.0 | 0.0 ± 0.0 | 0.0 ± 0.0 |
| 19 | TetraTetraTetra | 15.0 ± 0.0 | 17.2 ± 0.0 | 18.4 ± 0.0 | 10.9 ± 0.9 | 12.6 ± 0.5 | 9.5 ± 0.0 | 10.6 ± 0.0 |
| 20 | TetraTri(DAP)Anh I | 0.8 ± 0.0 | 0.0 ± 0.0 | 0.0 ± 0.0 | 0.8 ± 0.0 | 0.5 ± 0.0 | 0.0 ± 0.0 | 0.0 ± 0.0 |
| 20 | TetraTri(DAP)Anh II | 0.7 ± 0.0 | 0.0 ± 0.0 | 0.0 ± 0.0 | 1.4 ± 0.1 | 0.6 ± 0.0 | 0.0 ± 0.0 | 0.0 ± 0.0 |
| 21 | TetraTetraTetraTetra | 2.9 ± 0.0 | 4.4 ± 0.0 | 5.4 ± 0.1 | 1.1 ± 0.0 | 1.7 ± 0.0 | 0.9 ± 0.0 | 1.5 ± 0.0 |
| 22 | TetraTetraAnh I | 1.0 ± 0.0 | 2.3 ± 0.0 | 2.2 ± 0.0 | 1.9 ± 0.2 | 1.8 ± 0.0 | 2.9 ± 0.0 | 2.5 ± 0.0 |
| 23 | TeraTetraAnh II | 0.9 ± 0.0 | 1.7 ± 0.1 | 1.4 ± 0.0 | 1.7 ± 0.4 | 1.7 ± 0.0 | 3.5 ± 0.0 | 4.2 ± 0.0 |
| 24 | TetraTetraTetra Anh | 1.4 ± 0.0 | 3.3 ± 0.0 | 3.2 ± 0.0 | 1.5 ± 0.0 | 2.2 ± 0.0 | 2.9 ± 0.0 | 5.0 ± 0.0 |
| 25 | TetraTetraTetraTetra Anh | 0.5 ± 0.0 | 1.3 ± 0.0 | 1.3 ± 0.0 | 0.0 ± 0.0 | 0.5 ± 0.0 | 0.5 ± 0.0 | 1.4 ± 0.0 |
| % known peaks | | 88.1±0.2 | 94.3 ± 0.0 | 91.3 ± 0.1 | 90.5 ± 0.1 | 87.3 ± 0.2 | 95.4 ± 0.0 | 91.1 ± 0.0 |
| Monomers (Total) | | 24.0 ± 0.1 | 27.3 ± 0.0 | 25.0 ± 0.0 | 24.9 ± 7.1 | 24.5 ± 0.9 | 35.2 ± 0.0 | 32.8 ± 0.0 |
| Monomers di | | 0.0 ± 0.0 | 0.0 ± 0.0 | 0.0 ± 0.0 | 0.0 ± 0.0 | 0.0 ± 0.0 | 0.0 ± 0.0 | 0.3 ± 0.0 |
| Monomer tri | | 2.3 ± 0.0 | 0.1 ± 0.0 | 0.0 ± 0.0 | 2.3 ± 0.3 | 1.6 ± 0.1 | 0.7 ± 0.0 | 0.0 ± 0.0 |
| Monomer tetra | | 19.4 ± 0.1 | 27.0 ± 0.0 | 25.0 ± 0.0 | 21.0 ± 3.8 | 21.3 ± 1.5 | 34.1 ± 0.0 | 31.8 ± 0.0 |
| Monomer tetra-Gly4 | | 2.1 ± 0.0 | 0.1 ± 0.0 | 0.0 ± 0.0 | 1.3 ± 0.1 | 1.4 ± 0.0 | 0.0 ± 0.0 | 0.4 ± 0.0 |
| Monomer penta | | 0.2 ± 0.0 | 0.1 ± 0.0 | 0.0 ± 0.0 | 0.3 ± 0.0 | 0.2 ± 0.0 | 0.4 ± 0.0 | 0.3 ± 0.0 |
| Monomer anhydro | | 0.0 ± 0.0 | 0.3 ± 0.0 | 0.4 ± 0.0 | 0.0 ± 0.0 | 0.0 ± 0.0 | 0.0 ± 0.0 | 0.6 ± 0.0 |
| Dimers (Total) | | 50.3 ± 0.0 | 44.9 ± 0.0 | 43.9 ± 0.6 | 57.9 ± 1.6 | 53.6 ± 0.0 | 50.4 ± 0.0 | 46.8 ± 0.0 |
| Dimers (DD) | | 42.1 ± 0.1 | 44.4 ± 0.1 | 43.4 ± 0.8 | 47.2 ± 0.9 | 46.6 ± 0.1 | 50.1 ± 0.0 | 46.1 ± 0.0 |
| Dimers (LD) | | 8.2 ± 0.1 | 0.5 ± 0.0 | 0.5 ± 0.0 | 10.7 ± 0.1 | 7.0 ± 0.1 | 0.3 ± 0.0 | 0.7 ± 0.0 |
| Dimers anhydro | | 3.9 ± 0.0 | 4.3 ± 0.2 | 3.9 ± 0.0 | 6.3 ± 2.7 | 5.3 ± 0.1 | 6.7 ± 0.0 | 7.4 ± 0.0 |
| Trimers (Total) | | 21.8 ± 0.0 | 21.7 ± 0.0 | 23.7 ± 0.1 | 16.0 ± 1.6 | 19.3 ± 0.8 | 12.9 ± 0.0 | 17.1 ± 0.0 |
| Trimer anhydro | | 1.6 ± 0.0 | 3.5 ± 0.0 | 3.5 ± 0.0 | 1.6 ± 0.0 | 2.5 ± 0.0 | 3.0 ± 0.0 | 5.5 ± 0.0 |
| Tetramers (Total) | | 3.9 ± 0.0 | 6.0 ± 0.0 | 7.4 ± 0.2 | 1.2 ± 0.0 | 2.6 ± 0.0 | 1.4 ± 0.0 | 3.2 ± 0.0 |
| Tetramers (anhydro) | | 0.6 ± 0.0 | 1.3 ± 0.0 | 1.5 ± 0.0 | 0.0 ± 0.0 | 0.6 ± 0.0 | 0.5 ± 0.0 | 1.5 ± 0.0 |
| Dipeptides (Total) | | 0.0 ± 0.0 | 0.0 ± 0.0 | 0.0 ± 0.0 | 0.0 ± 0.0 | 0.0 ± 0.0 | 0.0 ± 0.0 | 0.3 ± 0.0 |
| Tripeptides (Total) | | 7.7 ± 0.0 | 0.4 ± 0.0 | 0.2 ± 0.0 | 8.9 ± 0.1 | 6.2 ± 0.0 | 0.8 ± 0.0 | 0.4 ± 0.0 |
| Tetrapeptides (Total) | | 92.0 ± 0.0 | 99.4 ± 0.0 | 99.7 ± 0.0 | 90.5 ± 0.0 | 93.6 ± 0.0 | 98.3 ± 0.0 | 98.7 ± 0.0 |
| Pentapeptides | | 0.3 ± 0.0 | 0.3 ± 0.0 | 0.1 ± 0.0 | 0.6 ± 0.0 | 0.2 ± 0.0 | 0.8 ± 0.0 | 0.6 ± 0.0 |
| 3-3-cross-linkage | | 4.6 ± 0.0 | 0.3 ± 0.0 | 0.2 ± 0.0 | 5.7 ± 0.0 | 3.9 ± 0.0 | 0.2 ± 0.0 | 0.4 ± 0.0 |
| 4-3-cross-linkage | | 38.1 ± 0.0 | 41.2 ± 0.0 | 43.1 ± 0.0 | 34.8 ± 2.1 | 37.7 ± 0.7 | 34.7 ± 0.0 | 36.9 ± 0.0 |
| Chain ends (anhydros) | | 2.6 ± 0.0 | 4.0 ± 0.0 | 3.9 ± 0.1 | 3.7 ± 0.8 | 3.6 ± 0.0 | 4.5 ± 0.0 | 6.5 ± 0.0 |
| Average chain length | | 37.8 ± 1.6 | 25.3 ± 1.7 | 26.0 ± 2.5 | 28.5 ± 46.2 | 27.5 ± 1.3 | 22.4 ± 0.0 | 15.4 ± 0.0 |
| Degree of cross-linkage | | 42.6 ± 0.0 | 41.5 ± 0.0 | 43.3 ± 0.0 | 40.5 ± 2.5 | 41.6 ± 0.5 | 34.9 ± 0.0 | 37.2 ± 0.0 |
| % peptides in cross-links | | 76.0 ± 0.1 | 72.7 ± 0.0 | 75.0 ± 0.0 | 75.1 ± 7.1 | 75.5 ± 0.9 | 64.8 ± 0.0 | 67.2 ± 0.0 |

**[a]Values are mean ± variation of two biological repeats.**

**Table S2:** Strains and plasmids used in this study

| Strain or Plasmid | Description | Reference or Source |
|---|---|---|
| **<u>Strains</u>** | | |
| *A. baumannii* AB5075 | Wild type | (1) |
| *A. baumannii* AB5075 | Δ*ldtJ* | (2) |
| *A. baumannii* AB5075 | Δ*mrcA* | (2) |
| *A. baumannii* AB5075 | Δ*ldtJ* Δ*mrcA* | This study |
| *A. baumannii* AB5075 | Δ*mrcB* | (2) |
| *A. baumannii* AB5075 | Δ*ldtJ* Δ*mrcB* | This study |
| *A. baumannii* AB5075 | Δ*ldtJ* Δ*mrcA*/pEV | This study |
| *A. baumannii* AB5075 | Δ*ldtJ* Δ*mrcA*/pLdtJ | This study |
| *A. baumannii* AB5075 | Δ*ldtJ* Δ*mrcA*/pPBP1A | This study |
| *A. baumannii* ATCC 17978 | Wild type | (3) |
| *A. baumannii* ATCC 17978 | Δ*ldtJ* | (4) |
| *A. baumannii* ATCC 17978 | Δ*mrcA* | (5) |
| *A. baumannii* ATCC 17978 | Δ*ldtJ* Δ*mrcA* | This study |
| *A. baumannii* ATCC 17978 | Δ*ldtJ* Δ*mrcA*/pEV | This study |
| *A. baumannii* ATCC 17978 | Δ*ldtJ* Δ*mrcA*/pLdtJ | This study |
| *A. baumannii* ATCC 17978 | Δ*ldtJ* Δ*mrcA*/pPBP1A | This study |
| *E. coli* MFD DAP- | Donor strain for conjugation | (6) |
| *E. coli* DH5α | Host strain for cloning | (7) |
| **<u>Plasmids</u>** | | |
| pABcre | Derived from pAT801-RA with *cre* recombinase and *arr2*, Rif$^{R}$ | (8, 9) |
| pJMP3665HygR | Replicative plasmid pJMP3665HygR, Hyg$^{R}$ | (10) |
| pJMP3665HygR*::mrcA* | Replicative plasmid pJMP3665HygR, Hyg$^{R}$ carrying AB5075 *mrcA* (*ABUW_0289*) | This study |
| pJMP3665HygR*::ldtJ* | Replicative plasmid pJMP3665HygR, Hyg$^{R}$ carrying AB5075 *ldtJ* (*ABUW_1189*) | This study |

| | | |
|---|---|---|
| pAT03 | pMMB67EH with FLP recombinase, Tet$^R$ | (11) |
| pAT04 | pMMB67EH with REC$_{Ab}$ system, Tet$^R$ | (11) |
| pKD4 | Kan$^R$ | (12) |
| pMMB67EH | pMMB67EH with the Kan$^R$ gene from pKD4 inserted into the PvuI site, Kan$^R$ | (4) |
| pABBR | pABBR_MCS with the Tet$^R$ gene from pKD4 replacing the *bla* gene, Tet$^R$ | (5) |
| pABBR*::mrcA* | pABBR, Tet$^R$ carrying 17978 *mrcA* (*A1S_3196-3197*) | (5) |
| pMMB67EH*::ldtJ* | pMMB67EH, Kan$^R$ carrying 17978 *ldtJ* (*A1S_2371*) | (4) |
| pMMB67EH*::mrcA-flag* | pMMB67EH, Kan$^R$ carrying the fusion 17978 *mrcA-flag* | This study |
| pMMB67EH*::ldtJ-flag* | pMMB67EH, Kan$^R$ carrying the fusion 17978 *ldtJ-flag* | This study |

**Table S3:** Primers used in this study.

| **Oligo Name** | **Sequence (5’ to 3’)** |
|---|---|
| **AB5075 Tn26 Insertion Primers** | |
| 5075 ABUW_1189 5’ | GCCCAACATTTCACTTACGC |
| 5075 ABUW_1189 3’ | CAATTGCATCTTTCCAGCAA |
| 5075 ABUW_0289 5’ | GGCGTTTCGCATTAATTGTT |
| 5075 ABUW_0289 3’ | ATATCAAAAGGCCGTTGCAG |
| 5075 ABUW_1358 5’ | CAGGTTCTACGCTGTCACCA |
| 5075 ABUW_1358 3’ | AATGCGCCAATTACTCAAGC |
| **ATCC 17978 Deletion Primers** | |
| A1S_2371 Kan-FRT 5’ | TTATATCCCTTCGCGTCTCAAATAAGCCAATATTAAATTCATAAGAATGAATGATTGGTGAGTTTATGGCCTAAAGGATCTGATTTTCCCTATTGCTTATATGAAAATTCTTAAGGTTGAATTAC**AGCGATTGTGTAGGCTGGAGCTGCTTCG** |
| A1S_2371 Kan-FRT 3’ | TTAGTAAACCTAGGCTGGTTTTATTTTTATAATCAAAACAATAACTACATATTCCACGGGGCTATGCTAAAAAATTTAATAAAAAAAGCCTGCATAAAGCAGGCTCTTTTAATTAAGAGGA**ATATCCTCCTTAGTTCCTATTCCG** |
| A1S_2371 confirm 5’ | TACTTGCAGCATGTTACATCGGGTTTA |
| A1S_2371 confirm 3’ | GGGTCAGATGCTGAAGCTGAATGGTTA |
| **Complementation Primers** | |
| 5075 *ldtJ* BamHI 5' | **CGCGGATCCATG**TTTGTTCGCTCATTACTCGCTATGAG |
| 5075 *ldtJ* XbaI 3' | **CGCTCTAGATTA**TTCTAAGAATTTAACAGTTACGCCTGAACG |
| 5075 *mrcA* BamHI 5' | **CGCGGATCCATG**AAAAAGCTATCCAGTTTGGGCTTCG |
| 5075 *mrcA* XbaI 3' | **CGCTCTAGATTA**TTCAATTTGATTAATCAAGTTTTCTAATTCATCTTTTTC |
| pJMP3665HygR confirm 5’ | GCAATTAATGTGAGTTAGCGCG |
| pJMP3665HygR confirm 3’ | CGTTCTGATTTAATCTGTATCAGGCT |
| **Co-immunoprecipitation Primers** | |
| 17978 *ldtJ-flag* KpnI 5’ | **CGCGGTACCATG**TTTGTTCGCTCATTACTCGC |
| 17978 *ldtJ-flag* BamHI 3’ | **CGCGGATCCTTATTTATCATCATCATCTTTATAATC**TTCTAAGAATTTAACAGTTACGCC |
| 17978 *mrcA-flag* KpnI 5’ | **CGCGGTACCATG**AAAAAGCTATCCAGTTTGGGCTTCG |

| | |
|---|---|
| | TGC |
| 17978 *mrcA-flag* BamHI 3’ | **CGCGGATCCTTATTTATCATCATCATCTTTATAATC**TTCAATTTGATTAATCAAGTTTTCTAATTCATCTTTTTCAC |
| pMMB67EH confirm 5’ | CGGTTCTGGCAAATATTCTGAAA |
| pMMB67EH confirm 3’ | CTGCGTTCTGATTTAATCTGTAT |
